# PFOS and GenX Induce Mitochondrial Toxicity and Differentially Alter Respiratory Complex Activities in Liver and Brain

**DOI:** 10.64898/2026.10.29.762821

**Authors:** Napissara Boonpraman, Da-Woon Kim, Shriya Pai, Nathan Kuhn, James O’Keefe, Shreesh Raj Sammi

## Abstract

Per- and polyfluoroalkyl substances (PFAS) are persistent environmental contaminants linked to adverse health effects, yet the mitochondrial consequences of exposure to individual PFAS and their mixtures remain incompletely understood. This study investigated the individual and combined effects of perfluorooctanesulfonic acid (PFOS) and hexafluoropropylene oxide dimer acid (GenX) on mitochondrial respiratory complex activities under acute, subchronic, and chronic exposure conditions. PFAS distribution in plasma, liver, and brain tissues, mitochondrial oxidative phosphorylation (OXPHOS) protein expression, and predicted interactions with mitochondrial Complexes III and IV were examined. Acute exposure inhibited Complex III and IV activities in isolated liver and brain mitochondria, with combined exposure generally producing greater inhibition than either compound alone. Following subchronic exposure (1 month), mitochondrial responses varied by dose and tissue type; liver enzymes showed significant inhibition of Complexes I to IV at 2 and 10 mg/kg BW, while brain mitochondria showed no alteration. After 5 months at 0.2 mg/kg BW, multiple liver complexes were inhibited, whereas the brain showed selective changes. PFOS and GenX were detected in plasma and brain tissues. In rats exposed to 2 mg/kg BW for 1 month, PFOS was higher than control in all three brain regions studied, with no difference between individual and combined exposure. OXPHOS protein expression varied across brain regions and exposure conditions. A comparison of internal levels in mice showed that co-exposure lowered GenX at 10 mg/kg BW, whereas at 0.2 mg/kg BW plasma PFOS and cortex GenX were higher with co-exposure. Plasma levels at 0.2 and 2 mg/kg BW were about ∼ 2.5 and 4.5 times the highest serum levels reported in the general and occupationally exposed populations, respectively. Molecular docking indicated distinct predicted interaction patterns, with PFOS generally exhibiting more favorable scores at Complex III binding sites and GenX at selected Complex IV regions. Overall, these findings demonstrate exposure-dependent and tissue-specific mitochondrial responses to PFOS and GenX and highlight the importance of evaluating combined PFAS exposure across multiple exposure durations.

## 1. Introduction

Per- and polyfluoroalkyl substances (PFAS) are a large group of synthetic chemicals extensively used in industrial processes and consumer products due to their chemical stability, thermal resistance, and water- and oil-repellent properties. Their widespread use and environmental persistence have resulted in extensive contamination of soil, air, food, and drinking water, raising concerns about long-term human exposure and associated health risks [1–3]. Among the various exposure pathways, contaminated drinking water represents an important source of human PFAS exposure [3].

Perfluorooctane sulfonate (PFOS), one of the most extensively studied legacy PFAS, is known for its environmental persistence, bioaccumulation, and prolonged retention in humans, particularly among occupationally exposed populations [4]. Although regulatory restrictions and manufacturing phase-outs have reduced environmental emissions and human exposure to certain legacy PFAS, these compounds remain widely detectable in the general population [5]. An analysis of the U.S. National Health and Nutrition Examination Survey (NHANES) reported a 16%–87% decline in geometric mean serum concentrations of four legacy PFAS between 1999–2000 and 2017–March 2020. Nevertheless, more than 96% of adolescents aged 12–19 years had measurable concentrations of these PFAS during 2017 to March 2020, indicating that exposure remains widespread despite reductions in legacy PFAS production [5]. In response to concerns about legacy PFAS, newer-generation alternatives have been introduced, including hexafluoropropylene oxide dimer acid (HFPO-DA, commonly known as GenX), which has been used as a replacement for perfluorooctanoic acid (PFOA) in certain industrial applications [6]. However, the introduction of replacement compounds has not eliminated existing legacy PFAS contamination. Both legacy and replacement PFAS have been detected in environmental waters, including drinking-water sources [6, 7]. Consequently, exposure to multiple PFAS may occur simultaneously, raising concerns about their potential combined toxicological effects.

PFOS has a long biological half-life (critically, the brain half-life is unknown) [8]. In a study on American blood donor samples, PFOS levels ranged from 0.0041 to 1.656 ppm [9]. Notably, studies in C. elegans have shown mitochondrial aberration at doses as low as 1 ppm [10]. Not only were these levels significantly higher than the Environmental Protection Agency’s current maximum contaminant level of 4 p.p.t. [11] (that used to be a 70 p.p.t. health advisory until the lower interim advisories of 2022 [12]), but they also suggest the bio-accumulative potential of PFAS. Salvatore et al. reported presumptive PFAS contamination sites in the U.S.A. to be ∼57,412 [13]. A recent study by Ruyle et al. encompassing 70% of the wastewater in the U.S.A. not only reported above-threshold PFAS contamination but also concluded that PFAS contamination is affecting drinking water sources for up to 23 million Americans [14]. PFAS is transported to organs through the blood, bound to serum albumin [15–17].

Understanding PFAS-associated toxicity requires consideration of exposure concentration, duration, and toxicokinetic behavior, as absorption, distribution, and elimination vary among PFAS and across species. Studies of PFOS elimination have reported substantial differences in serum half-lives across species, with retention lasting several years in humans compared with weeks to months in rodents and nonhuman primates [4, 8]. Studies in Sprague–Dawley rats have also demonstrated tissue-dependent distribution and isomer-specific elimination of PFOS following single oral and repeated dietary exposure [9, 10]. In contrast, GenX is eliminated more rapidly in rodents, primarily through urinary excretion [21]. Paradoxically, while a significant level of GenX was present in water, it was not detectable in humans in a North Carolina biomonitoring study. This does not indicate a subdued risk, but was due to the consumption of bottled water months earlier, when samples were taken [22], and a shorter half-life [23], implying that GenX can circumvent detection.[11] These differences in toxicokinetics may influence internal PFAS concentrations and biological responses during repeated exposure and should be considered when extrapolating findings from animal models to humans [1]. Evaluating different exposure concentrations and durations may help assess the toxicological responses to individual PFAS and their mixtures.

The liver is a major target organ of PFAS toxicity, with experimental studies reporting hepatic accumulation and disturbances in lipid metabolism [1, 12]. PFAS exposure has also been associated with alterations in peroxisome proliferator-activated receptor α (PPARα)-related signaling, which plays an important role in hepatic lipid metabolism [12]. Although GenX has a shorter biological half-life than PFOS, experimental studies have reported increased liver weight, histopathological changes, and altered hepatic lipid metabolism following GenX exposure in mice [13, 14]. In human HepG2 cells, GenX has also been shown to induce reactive oxygen species (ROS) generation and apoptosis, accompanied by changes in apoptosis-related proteins [15].

Although the brain is protected by the blood–brain barrier (BBB), several PFAS have been detected in human brain tissues, with distribution patterns varying among individual compounds [16]. More recent analyses of paired human plasma and brain tissue samples have identified compound-dependent differences in brain-to-plasma concentration ratios, providing further evidence that certain PFAS can cross the BBB [17]. Experimental studies have also reported PFOS-induced apoptosis in cerebellar granule cells through reactive oxygen species (ROS)-dependent signaling and increased susceptibility of dopaminergic terminals to subsequent neurotoxic insult [18, 19]. GenX has been shown to interfere with BBB efflux transporter activity, while chronic exposure in mice has been associated with brain accumulation, BBB dysfunction, neuroinflammation, and behavioral alterations [20, 21]. These findings suggest that both legacy and replacement PFAS can affect the central nervous system. However, the effects of exposure concentration and duration on PFOS and/or GenX distribution among different brain regions remain unclear.

Mitochondrial dysfunction has been implicated in PFAS-associated toxicity in multiple organs, including the liver and brain. Mitochondria are essential for cellular energy production and metabolic regulation, with neurons being particularly dependent on oxidative phosphorylation to maintain energy-demanding functions [22]. Mitochondrial respiratory complexes I–IV facilitate electron transport, while proton translocation by Complexes I, III, and IV contributes to the electrochemical gradient required for ATP synthesis [23, 24]. Previous studies have reported that PFAS can alter mitochondrial respiration, membrane function, and oxidative balance, with effects varying according to chemical structure, exposure concentration, and experimental model [25, 26]. PFOS exposure has been shown to increase ROS generation and reduce mitochondrial content in *Caenorhabditis elegans*, as well as impair mitochondrial respiration in human SH-SY5Y cells [27]. GenX exposure has been linked to mitochondrial dysfunction and inflammatory responses in astrocytes, as well as ROS generation and apoptosis in primary cortical neurons [28, 29]. Previous studies from our group have further demonstrated that PFOS inhibits mitochondrial respiratory complexes II–IV, while glutathione (GSH) and γ-glutamyl-cysteine (γ-Glu-Cys) can alleviate Complex III inhibition [30]. In addition, a comparative study of legacy and replacement PFAS reported compound-dependent inhibition of respiratory complexes I–IV in isolated rat brain mitochondria, with GenX primarily affecting Complex IV activity [31]. However, mitochondrial responses are not consistent across experimental models, as GenX did not significantly alter oxidative phosphorylation or ATP levels in zebrafish embryos and larvae under the conditions tested [32].

Previous studies of PFAS mixtures have reported combined cytotoxic and neurotoxic effects, including responses consistent with concentration addition at low effect levels [33]. PFAS mixture exposure has also been associated with mitochondrial dysfunction and alterations in glucose and lipid metabolism in zebrafish larvae [34]. However, findings from different PFAS mixtures and experimental models may not directly apply to combined PFOS and GenX exposure in the mammalian liver and brain. In particular, it remains unclear how mitochondrial respiratory complex activities respond to combined PFOS and GenX exposure compared with either compound alone, and how these responses vary across tissues, exposure concentrations, and durations.

In this study, we aimed to investigate the individual and combined effects of PFOS and GenX under acute, sub-chronic, and chronic exposure conditions, focusing on mitochondrial respiratory complex activities in the liver and brain. We also examined OXPHOS protein expression across different brain regions and measured PFOS and GenX concentrations in plasma and tissues to characterize their distribution following repeated exposure.

## 2. Materials and Methods

### 2.1. Animals and chemical exposure

Adult male and female C57BL/6J mice were obtained from The Jackson Laboratory (strain #000664). Mice were approximately 12 weeks old and weighed approximately 28–30 g at the start of the experiments. Animals were acclimated for at least 7 days before treatment. All animal procedures were approved by the Institutional Animal Care and Use Committee (IACUC) at Michigan State University and were conducted in accordance with institutional guidelines.

For the acute treatment experiments, 3-month-old male and female mice were used (n = 3 per treatment group per sex).

For the subchronic and chronic PFOS and GenX exposure studies, male mice were assigned to four treatment groups: vehicle control (0.5% Tween 20), PFOS, GenX, or combined PFOS and GenX. For the 30-day exposure study, mice received PFOS, GenX, or PFOS + GenX at 0.2, 2, or 10 mg/kg body weight (BW)/day (n = 12 per group). In the combined exposure groups, each compound was administered at the corresponding dose. For the chronic exposure study, mice received PFOS, GenX, or PFOS + GenX at 0.2 mg/kg BW/day for 5 months (n = 10 per group).

All repeated exposures were administered *ad libitum* through the drinking water. Body weight and water consumption were monitored at least twice weekly. Treatment solutions were prepared weekly, and chemical concentrations were adjusted based on measured body weight and water consumption to maintain the intended daily doses.

Adult Fischer (CDF) rats (F344/DuCrl, strain 002) were obtained from Charles River Laboratories (Wilmington, MA, USA) or bred in-house at Michigan State University. Two age groups were included: 6-month-old rats weighing approximately 350–380 g and 10-month-old rats weighing approximately 360–400 g at the start of treatment. Animals were acclimated for at least 7 days before treatment.

For the acute treatment experiments, Male and female rats were used (n = 4 per treatment group per sex). For the repeated-exposure study, rats were assigned to four treatment conditions: vehicle control (0.5% Tween 20), PFOS, GenX, or P+G. Rats received PFOS, GenX, or P+G at 2 mg/kg BW/day for 30 days through the drinking water ad libitum (n = 6 per group).

### 2.2. Tissue collection and sample preparation

At the end of the treatment period, mice were euthanized by cervical dislocation, whereas rats were euthanized by decapitation. Immediately following euthanasia, blood was collected directly from the heart into K_2_EDTA-containing tubes. Blood samples were centrifuged at 1,500 × *g* for 10 min at 4°C, and the resulting plasma was collected and stored at −80°C until analysis of PFOS and GenX concentrations by liquid chromatography–tandem mass spectrometry (LC–MS/MS).

Brain and liver tissues were rapidly collected and allocated for different experimental analyses. Brain tissues were used for Western blotting (WB), immunohistochemistry (IHC), LC–MS/MS, and mitochondrial enzyme activity (EA) assays. Brain regions collected for biochemical analyses included the cortex, hippocampus, cerebellum, and brainstem. Fresh brain and liver tissues designated for mitochondrial analysis were processed immediately for mitochondrial isolation, whereas samples designated for WB or LC–MS/MS analysis were stored at −80°C until further analysis. Tissues designated for histological analyses were processed separately as described below.

Tissues were allocated among the different experimental endpoints so that each analysis included samples from six individual animals (n = 6 per treatment group). For studies requiring separate tissue preparations, animals were divided into sets according to the downstream analysis. One set was used primarily for WB and LC–MS/MS analyses, while a separate set was used for mitochondrial isolation and histological analysis.

### 2.3. Mitochondrial isolation

Mitochondria were isolated from brain and liver tissues by differential centrifugation using a modified protocol based on Frezza et al. (2007) [35]. Briefly, freshly collected tissues were homogenized in ice-cold mitochondrial isolation buffer, followed by differential centrifugation to obtain mitochondrial fractions. The isolated mitochondria were washed, resuspended in isolation buffer, and stored at −80°C until further analysis. Mitochondrial protein concentrations were determined using the BCA assay prior to subsequent experiments.

### 2.4. Mitochondrial Respiratory Chain Enzyme Activity Assays

Mitochondrial respiratory chain (RC) enzyme activities were measured in isolated mitochondrial fractions from brain and liver tissues using spectrophotometric assays, as previously described [31, 36, 37]. The activities of respiratory chain complexes I–IV were normalized to citrate synthase (CS) activity to account for differences in mitochondrial content.

Complex I (NADH oxidoreductase) activity was determined by monitoring NADH oxidation at 340 nm. Complex II (succinate dehydrogenase) activity was measured by monitoring the reduction of 2,6-dichlorophenolindophenol (DCPIP) at 600 nm. Complex III (ubiquinol c oxidoreductase) activity was assessed by monitoring cytochrome c reduction at 550 nm, whereas Complex IV (cytochrome c oxidase) activity was measured by monitoring the oxidation of reduced cytochrome c at 550 nm. Following substrate addition, absorbance was recorded every 10 s for 3 min. Enzyme activities were normalized to the corresponding CS activity.

For the acute *ex vivo* exposure experiments, mitochondria were isolated from the brain and liver tissues of untreated 3-month-old mice and 6-month-old rats. Following isolation, mitochondrial fractions were directly exposed to PFOS, GenX, or a combination of PFOS and GenX (P+G) for 15 min. Mitochondrial respiratory chain complex I–IV activities were subsequently measured to evaluate the acute effects of individual and combined PFAS exposure on mitochondrial function.

### 2.5. Western blots

Protein concentrations were measured with the Pierce™ BCA Protein Assay Kit, and equal amounts of protein (24 µg per sample) were separated on 4–20% Mini-PROTEAN TGX Precast Protein Gels and transferred onto 0.2 µm nitrocellulose membrane. Membranes were blocked in 5% BSA and probed with primary antibodies against total OXPHOS (ab110413) and Hsp60 (ab190828), followed by IRDye 680RD and IRDye 800CW secondary antibodies. Blots were scanned on an Odyssey CLx and quantified using Image Studio (version 6.1). Full details are in the Supplementary Information (Section S6).

### 2.6. LC-MS/MS

Plasma, brain, and liver samples were flash-frozen and stored at −80 °C until PFOS and GenX quantification by LC–MS/MS at the Michigan State University Mass Spectrometry and Metabolomics Core Facility.

Samples were extracted using 75% methanol containing 13C-labeled internal standards. An aliquot (1 µL) of each extract was injected into a Waters BEH C18 column (2.1 × 50 mm) coupled to a Waters ACQUITY UPLC system and Xevo TQ-XS triple quadrupole mass spectrometer. Chromatographic separation was performed using 2 mM ammonium acetate in water (mobile phase A) and 2 mM ammonium acetate in methanol (mobile phase B), with a gradient from 5% to 95% B over 5 min, followed by re-equilibration to initial conditions. The flow rate was 0.3 mL/min, and the column temperature was maintained at 35 °C. Analytes were detected using negative electrospray ionization (ESI) with compound-specific MS/MS parameters. Data were processed using TargetLynx in MassLynx software (Waters Corporation), and tissue PFAS concentrations were normalized to sample protein content.

### 2.7. Chemicals

Perfluorooctanesulfonic acid potassium salt (PFOS; Cat. #0612) was purchased from AK Scientific (Union City, CA, USA). Ammonium perfluoro(2-methyl-3-oxahexanoate) (GenX; Cat. #C23990) was obtained from Manchester Organics (Cheshire, UK).

### 2.8. Molecular Docking

Molecular docking was performed to evaluate the potential interactions of PFOS, GenX, and their combination with mitochondrial respiratory Complexes III and IV in human and mouse protein structures. Protein structures were obtained from the Protein Data Bank (PDB IDs: 9HZL for human Complex III, 7O3C for mouse Complexes III and IV, and 5Z62 for human Complex IV). The protein structures were processed using Biopython and PDBFixer, then converted to PDBQT format using ADFR Suite. Important cofactors, including hemes, iron–sulfur clusters, and copper centers, were retained. PFOS and GenX were prepared in their negatively charged forms using RDKit and Meeko. The Qo and Qi sites of Complex III were identified using the experimentally resolved ubiquinone molecules in the human structure, and the equivalent mouse sites were identified by structural alignment. For Complex IV, the heme a₃–CuB catalytic region was used as the docking region. All calculations were performed using AutoDock Vina v1.2.7 with a rigid receptor. PFOS and GenX were docked individually and simultaneously, with the mixture represented by one molecule of each compound. Full details are in the Supplementary Information (Section S7).

### 2.9. Statistical Analysis

Statistical analyses were performed using GraphPad Prism (version 10.6.1; GraphPad Software, Inc., La Jolla, CA, USA). Data are presented as mean ± standard error of the mean (SEM). Each experiment included at least three independent biological replicates. Comparisons between two groups were performed using unpaired Welch’s *t*-test, whereas comparisons among multiple groups were analyzed using one-way or two-way analysis of variance (ANOVA), followed by appropriate post hoc multiple-comparison tests, as specified in the figure legends. A *p*-value < 0.05 was considered statistically significant.

## 3. Results

### 3.1. Combined exposure to PFOS and GenX produces greater inhibition of mitochondrial Complex III and IV activities in the liver

#### 3.1.1. Acute exposure to PFOS and GenX inhibits mitochondrial Complex III activity in the liver

In the present study, we examined the effects of PFOS and GenX, individually and in combination, on mitochondrial respiratory complex activities in isolated mouse liver mitochondria. Isolated mouse liver mitochondria were exposed to PFOS, GenX, or their combination (P+G) at 25, 50, and 75 ppm for 15 min. The experiments included biological replicates from male and female mice (at least n = 3–6 per group). Data are presented as mean ± SEM. Detailed values for each concentration are in the Supplementary Information (Section S1).

PFOS significantly reduced Complex III activity at all concentrations tested (control, 1.000 ± 0.000; 0.789 ± 0.036, p = 0.0205 at 25 ppm; 0.677 ± 0.054, p = 0.0007 at 50 ppm; and 0.607 ± 0.052, p < 0.0001 at 75 ppm) compared with the control (Fig. 1A). GenX also reduced Complex III activity, but not significantly at 25 ppm; significant reductions were observed at 50 ppm (0.798 ± 0.017, p = 0.0234) and 75 ppm (0.607 ± 0.022, p = 0.0002) (Fig. 1B). Combined exposure to PFOS and GenX (P+G) significantly reduced Complex III activity at all concentrations (0.653 ± 0.066, p = 0.0007 at 25 ppm; 0.370 ± 0.058, p < 0.0001 at 50 ppm; and 0.314 ± 0.067, p < 0.0001 at 75 ppm) (Fig. 1C). When the treatments were compared at the same concentration (Fig. 1D–F), no significant differences were detected at 25 ppm, whereas the P+G group showed significantly lower activity than PFOS alone and GenX alone at 50 ppm (0.370 ± 0.058 vs. 0.677 ± 0.054, p = 0.0008; and vs. 0.798 ± 0.017, p < 0.0001) and 75 ppm (0.314 ± 0.067 vs. 0.607 ± 0.052, p = 0.0055; and vs. 0.607 ± 0.022, p = 0.0082). Antimycin A, a Complex III inhibitor, significantly reduced activity (0.237 ± 0.035, p < 0.0001) (Fig. 1G).

**Figure 1.**
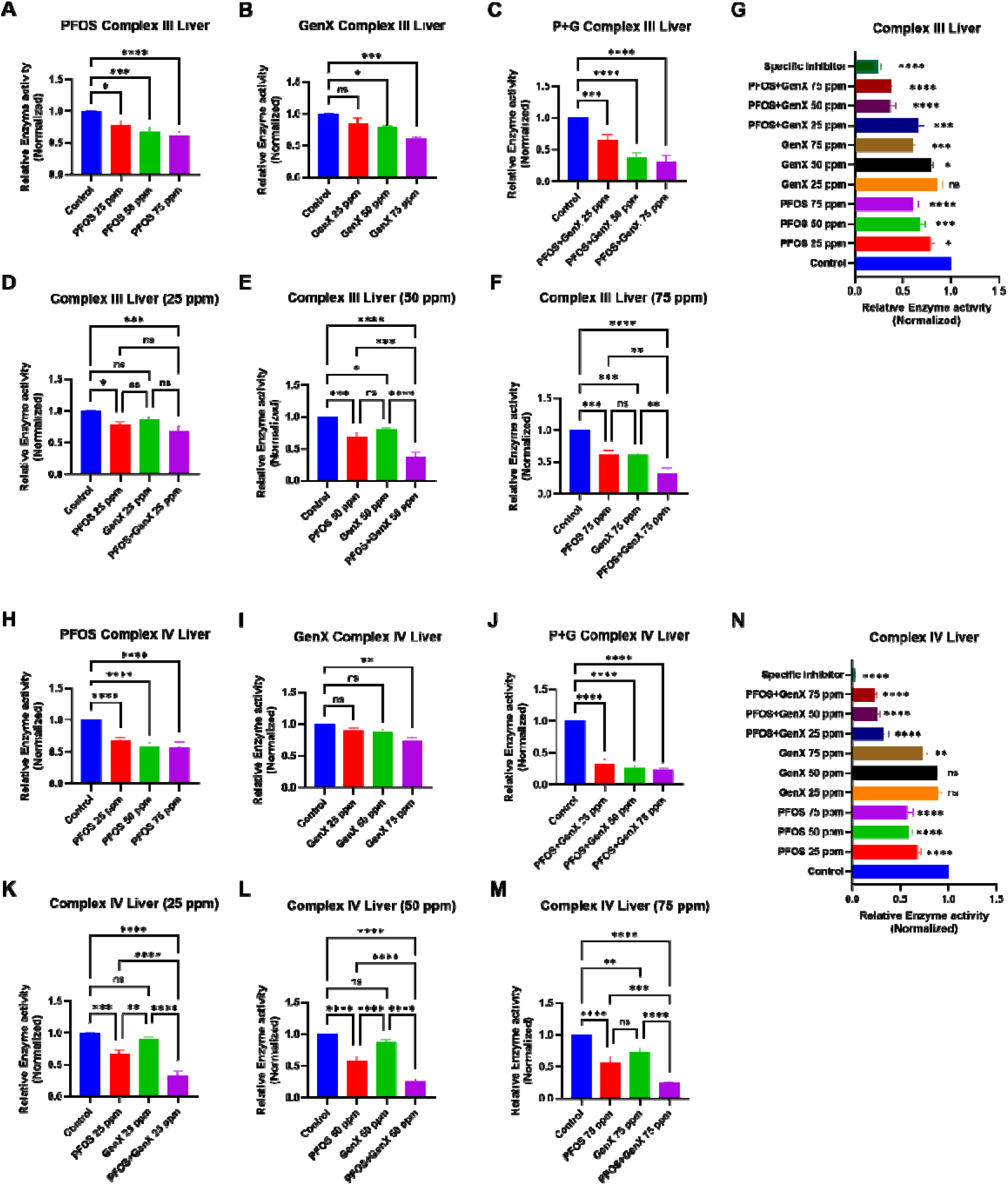
Effects of acute PFOS and GenX exposure on mitochondrial Complex III and IV activities in isolated mouse liver mitochondria. Isolated mouse liver mitochondria were exposed to PFOS, GenX, or their combination (P+G) at 25, 50, and 75 ppm for 15 min. (A–C) Complex III activities following exposure to (A) PFOS, (B) GenX, and (C) P+G at different concentrations. (D–F) Comparisons of Complex III activities among PFOS, GenX, and P+G treatments at (D) 25 ppm, (E) 50 ppm, and (F) 75 ppm. (G) Comparison of Complex III activities across individual and combined treatments, including a Complex III-specific inhibitor control (antimycin A). (H–J) Complex IV activities following exposure to (H) PFOS, (I) GenX, and (J) P+G at different concentrations. (K–M) Comparisons of Complex IV activities among PFOS, GenX, and P+G treatments at (K) 25 ppm, (L) 50 ppm, and (M) 75 ppm. (N) Complex IV activity following treatment with potassium cyanide (KCN), a Complex IV-specific inhibitor. Enzyme activities were normalized to the control group. Data are presented as mean ± SEM (n = 3–6 independent biological replicates per group from both male and female mice). Statistical significance was determined using one-way ANOVA followed by Dunnett’s multiple-comparisons test for panels A–C, G, H–J, and N, and Tukey’s multiple-comparisons test for panels D–F and K–M. \**p* < 0.05, \*\**p* < 0.01, \*\*\**p* < 0.001, \*\*\*\**p* < 0.0001; ns, not significant.

Both PFOS and GenX inhibited mitochondrial Complex III activity following acute exposure, although PFOS produced a significant reduction at a lower concentration than GenX. Combined PFOS and GenX exposure produced greater inhibition of Complex III activity than either compound alone at 50 and 75 ppm, whereas no significant differences among treatment groups were observed at 25 ppm.

#### 3.1.2. Acute exposure to PFOS and GenX inhibits mitochondrial Complex IV activity in the liver

PFOS significantly reduced Complex IV activity at all concentrations tested (control, 1.000 ± 0.000; 0.675 ± 0.040, p < 0.0001 at 25 ppm; 0.578 ± 0.048, p < 0.0001 at 50 ppm; and 0.569 ± 0.064, p < 0.0001 at 75 ppm) (Fig. 1H). GenX did not significantly alter Complex IV activity at 25 or 50 ppm, but significantly reduced activity at 75 ppm (0.732 ± 0.046, p = 0.0047) (Fig. 1I). P+G significantly reduced Complex IV activity at all concentrations (0.326 ± 0.054, p < 0.0001 at 25 ppm; 0.260 ± 0.025, p < 0.0001 at 50 ppm; and 0.236 ± 0.019, p < 0.0001 at 75 ppm) (Fig. 1J). At each concentration, the P+G group showed significantly lower Complex IV activity than either PFOS or GenX alone (0.326 ± 0.054, p < 0.0001 versus both at 25 ppm; 0.260 ± 0.025, p < 0.0001 versus both at 50 ppm; and 0.236 ± 0.019, p = 0.0006 versus PFOS and p < 0.0001 versus GenX at 75 ppm) (Fig. 1K–M). PFOS also showed significantly lower activity than GenX at 25 ppm (0.675 ± 0.040 vs. 0.897 ± 0.033, p = 0.0056) and 50 ppm (0.578 ± 0.048 vs. 0.877 ± 0.025, p < 0.0001), but not at 75 ppm. Potassium cyanide (KCN), a Complex IV inhibitor, significantly reduced activity (0.025 ± 0.001, p < 0.0001) (Fig. 1N).

Similar to the findings for Complex III, both PFOS and GenX inhibited Complex IV activity following acute exposure, although their effects differed. PFOS significantly reduced Complex IV activity at all concentrations, whereas GenX showed significant inhibition only at 75 ppm. Combined exposure to PFOS and GenX significantly reduced Complex IV activity compared with either compound alone at all concentrations tested. In contrast, significant differences between combined and individual treatments were observed for Complex III only at 50 and 75 ppm. This difference suggests that the effects of combined PFAS exposure vary between mitochondrial respiratory complexes.

#### 3.1.3. Acute exposure to PFOS and GenX inhibits mitochondrial Complex III activity in the brain

To determine whether PFOS and GenX also affect mitochondrial respiratory function in the brain, isolated brain mitochondria were exposed to PFOS, GenX, or their combination (P+G) for 15 min.

In brain mitochondria, PFOS significantly reduced Complex III activity at all concentrations tested (control, 1.000 ± 0.000; 0.801 ± 0.052, p = 0.0068 at 25 ppm; 0.737 ± 0.011, p = 0.0006 at 50 ppm; and 0.662 ± 0.050, p < 0.0001 at 75 ppm) (Fig. 2A). GenX did not significantly affect Complex III activity at 25 ppm, but significantly reduced activity at 50 ppm (0.892 ± 0.063, p = 0.0163) and 75 ppm (0.870 ± 0.048, p = 0.0013) (Fig. 2B). P+G significantly reduced Complex III activity at all concentrations (0.767 ± 0.066, p = 0.0042 at 25 ppm; 0.610 ± 0.067, p < 0.0001 at 50 ppm; and 0.441 ± 0.065, p < 0.0001 at 75 ppm) (Fig. 2C). PFOS produced significantly lower activity than GenX at 25, 50, and 75 ppm (0.801 ± 0.052 vs. 1.007 ± 0.020, p = 0.0162; 0.737 ± 0.011 vs. 0.892 ± 0.063, p = 0.0144; and 0.662 ± 0.050 vs. 0.870 ± 0.048, p = 0.0012). The P+G group showed significantly lower activity than GenX at all three concentrations (0.767 ± 0.066 vs. 1.007 ± 0.020, p = 0.0051 at 25 ppm; 0.610 ± 0.067 vs. 0.892 ± 0.063, p < 0.0001 at 50 ppm; and 0.441 ± 0.065 vs. 0.870 ± 0.048, p < 0.0001 at 75 ppm) and than PFOS at 50 ppm (0.610 ± 0.067 vs. 0.737 ± 0.011, p = 0.0496) and 75 ppm (0.441 ± 0.065 vs. 0.662 ± 0.050, p = 0.0006) (Fig. 2D–F). Antimycin A significantly reduced Complex III activity (0.086 ± 0.017, p < 0.0001) (Fig. 2G).

**Figure 2.**
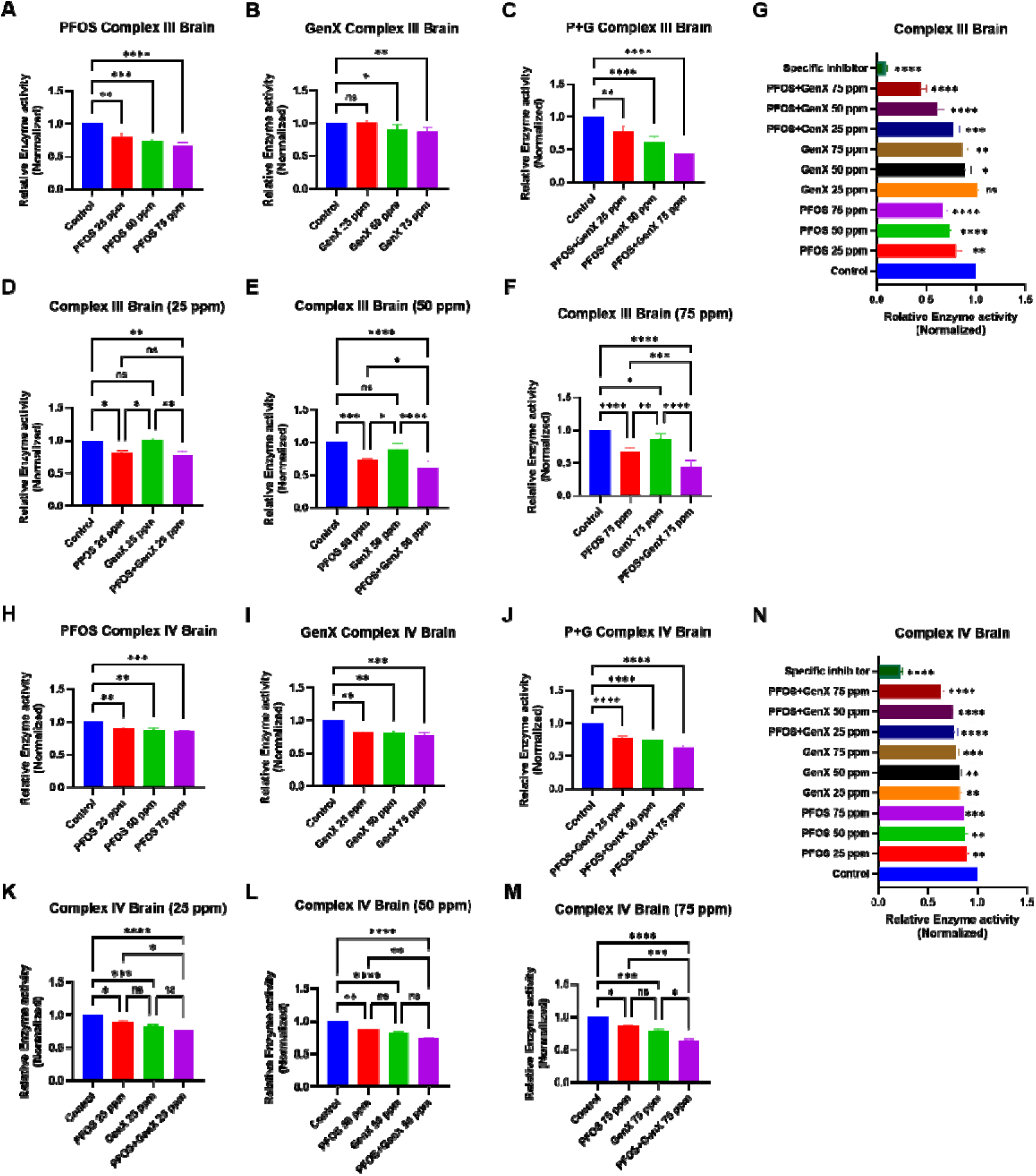
Effects of acute PFOS and GenX exposure on mitochondrial Complex III and IV activities in isolated mouse brain mitochondria. Isolated mouse brain mitochondria were exposed to PFOS, GenX, or their combination (P+G) at 25, 50, and 75 ppm for 15 min. (A–C) Complex III activities following exposure to (A) PFOS, (B) GenX, and (C) P+G at different concentrations. (D–F) Comparisons of Complex III activities among PFOS, GenX, and P+G treatments at (D) 25 ppm, (E) 50 ppm, and (F) 75 ppm. (G) Comparison of Complex III activities across individual and combined treatments, including antimycin A as a Complex III-specific inhibitor control. (H–J) Complex IV activities following exposure to (H) PFOS, (I) GenX, and (J) P+G at different concentrations. (K–M) Comparisons of Complex IV activities among PFOS, GenX, and P+G treatments at (K) 25 ppm, (L) 50 ppm, and (M) 75 ppm. (N) Complex IV activity following treatment with potassium cyanide (KCN) as a Complex IV-specific inhibitor control. Enzyme activities were normalized to the control group. Data are presented as mean ± SEM (n = 3–6 independent biological replicates per group from both male and female mice). Statistical significance was determined using one-way ANOVA followed by Dunnett’s multiple-comparisons test for panels A–C, G, H–J, and N, and Tukey’s multiple-comparisons test for panels D–F and K–M. *p < 0.05, **p < 0.01, ***p < 0.001, ****p < 0.0001; ns, not significant.

Both PFOS and GenX inhibited mitochondrial Complex III activity following acute exposure, although PFOS produced a significant reduction at a lower concentration than GenX. This pattern is consistent with the response observed in liver mitochondria, where PFOS also inhibited Complex III activity at lower concentrations than GenX. However, the effects of combined exposure differed between the two tissues. In brain mitochondria, P+G produced significantly greater inhibition than GenX at all concentrations, whereas the difference between P+G and PFOS was significant only at 50 and 75 ppm. These findings suggest that PFOS contributes substantially to the inhibition of brain mitochondrial Complex III at lower concentrations, while combined exposure produces greater inhibition at higher concentrations.

#### 3.1.4. Acute exposure to PFOS and GenX inhibits mitochondrial Complex IV activity in the brain

PFOS (control, 1.000 ± 0.000; 0.888 ± 0.020, p = 0.0052 at 25 ppm; 0.872 ± 0.024, p = 0.0017 at 50 ppm; and 0.861 ± 0.001, p = 0.0008 at 75 ppm), GenX (0.818 ± 0.024, p = 0.0040 at 25 ppm; 0.816 ± 0.015, p = 0.0035 at 50 ppm; and 0.781 ± 0.027, p = 0.0008 at 75 ppm), and P+G (0.770 ± 0.042, p < 0.0001 at 25 ppm; 0.744 ± 0.002, p < 0.0001 at 50 ppm; and 0.630 ± 0.026, p < 0.0001 at 75 ppm) each significantly reduced brain Complex IV activity at all concentrations tested compared with the control (Fig. 2H–J). The P+G group showed significantly lower Complex IV activity than PFOS alone at 25 ppm (0.770 ± 0.042 vs. 0.888 ± 0.020, p = 0.0261), 50 ppm (0.744 ± 0.002 vs. 0.872 ± 0.024, p = 0.0031), and 75 ppm (0.630 ± 0.026 vs. 0.861 ± 0.001, p = 0.0004), and than GenX alone at 75 ppm (0.630 ± 0.026 vs. 0.781 ± 0.027, p = 0.0015) (Fig. 2K–M). No significant difference was observed between PFOS and GenX at any concentration. KCN significantly reduced Complex IV activity (0.225 ± 0.018, p < 0.0001) (Fig. 2N).

In contrast with Complex III, Complex IV activity was significantly reduced by both PFOS and GenX at all concentrations tested in brain mitochondria. This differs from the liver mitochondrial response, where GenX significantly inhibited Complex IV only at 75 ppm. The results suggest that GenX affects mitochondrial respiratory complexes differently in liver and brain mitochondria, although direct statistical comparisons between tissues would be needed to establish differences in sensitivity.

### 3.2. Subchronic and chronic PFOS and GenX exposure alters mitochondrial complex activities in the liver

To examine the effects of repeated PFAS exposure on mitochondrial complex activities, mice were treated with PFOS, GenX, or their combination (P+G) at 0.2, 2, and 10 mg/kg BW for 1 month. An additional group received 0.2 mg/kg BW for 5 months to examine the effects of longer-term exposure. Mitochondrial Complex I–IV enzyme activities were measured in isolated liver mitochondria and normalized to citrate synthase (CS) activity. Data are presented as mean ± SEM (Table S1). Detailed values are in the Supplementary Information (Section S2, Tables S1 and S2).

The 1-month results show that liver mitochondrial respiratory enzyme activities were affected differently across exposure levels. At 0.2 mg/kg BW, combined PFOS and GenX exposure significantly increased liver mitochondrial Complex I activity after 1 month (control, 0.933 ± 0.099; 1.360 ± 0.064, p = 0.0391), whereas no significant changes were observed in Complex II–IV activities (Fig. 3A). In contrast, at 2 and 10 mg/kg BW, several respiratory complexes showed significantly reduced activities. At 2 mg/kg BW, Complex I and II activities were significantly lower in the GenX and P+G groups (Complex I, 0.563 ± 0.032, p = 0.0017 and 0.532 ± 0.054, p = 0.0007; Complex II, 0.459 ± 0.047, p < 0.0001 and 0.464 ± 0.095, p = 0.0001), Complex III activity in all treatment groups (0.662 ± 0.112, p = 0.0189; 0.471 ± 0.043, p = 0.0001; and 0.354 ± 0.068, p < 0.0001, for PFOS, GenX, and P+G, respectively), and Complex IV activity in the GenX and P+G groups (0.468 ± 0.029, p = 0.0001 and 0.449 ± 0.076, p < 0.0001) compared with the control (Complex I, 1.000 ± 0.078; Complex II, 1.000 ± 0.146; Complex III, 1.000 ± 0.116; Complex IV, 1.000 ± 0.034) (Fig. 3B). At 10 mg/kg BW, all three treatments significantly reduced Complex I–IV activities compared with the control (Complex I: control, 0.979 ± 0.050; PFOS, 0.697 ± 0.050, p = 0.0007; GenX, 0.437 ± 0.079, p < 0.0001; P+G, 0.513 ± 0.049, p < 0.0001. Complex II: control, 0.994 ± 0.022; PFOS, 0.483 ± 0.068; GenX, 0.184 ± 0.032; P+G, 0.322 ± 0.020; p < 0.0001 for all. Complex III: control, 1.000 ± 0.096; PFOS, 0.638 ± 0.042; GenX, 0.343 ± 0.035; P+G, 0.423 ± 0.014; p < 0.0001 for all. Complex IV: control, 1.009 ± 0.033; PFOS, 0.546 ± 0.028; GenX, 0.344 ± 0.096; P+G, 0.345 ± 0.044; p < 0.0001 for all) (Fig. 3C).

**Figure 3.**
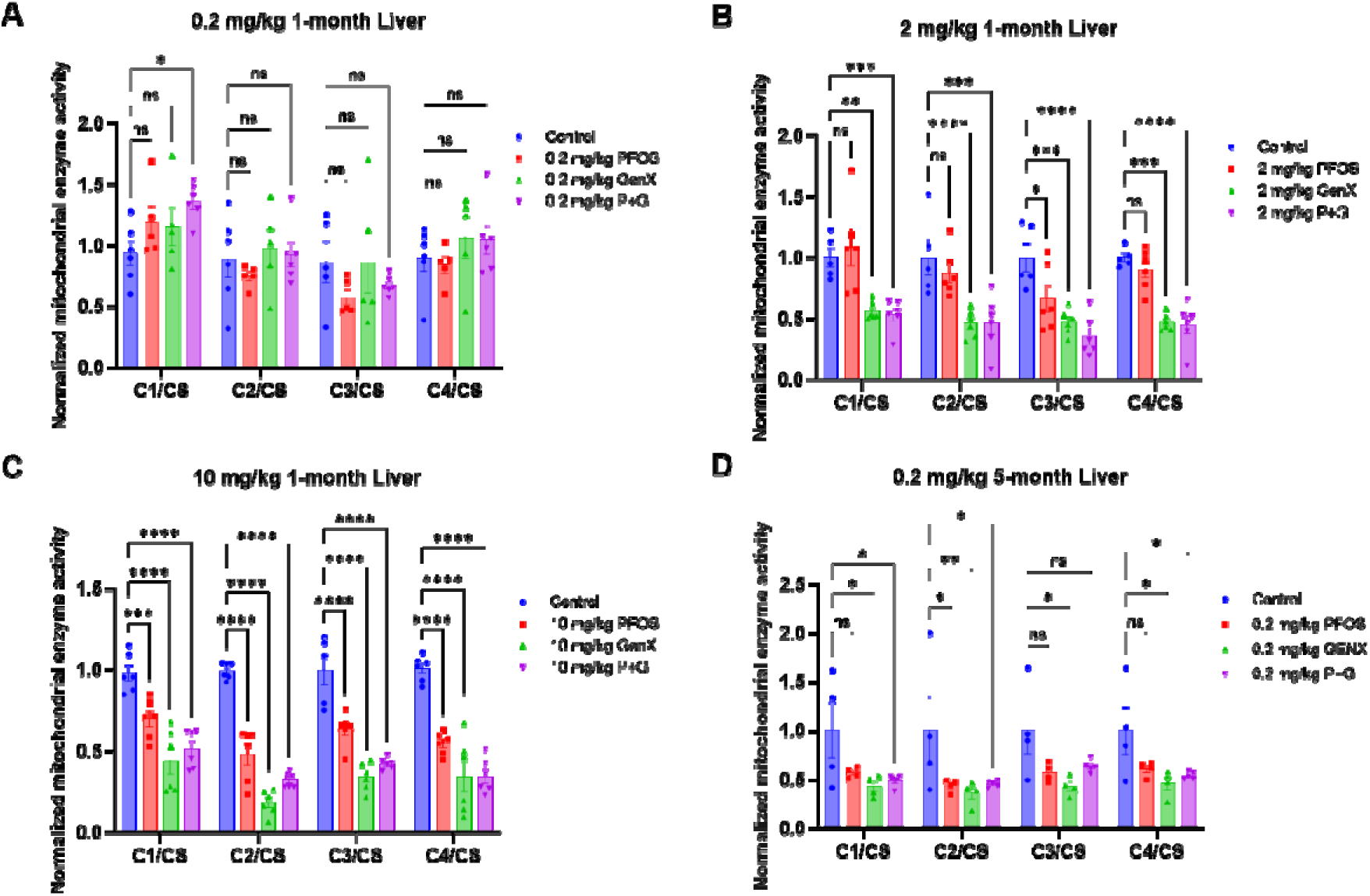
Effects of sub-chronic and chronic PFOS and GenX exposure on mitochondrial respiratory complex activities in mouse liver mitochondria. Mice were exposed to PFOS, GenX, or their combination (P+G) at 0.2, 2, and 10 mg/kg BW for 1 month or at 0.2 mg/kg BW for 5 months. Mitochondrial Complex I–IV activities were measured in isolated liver mitochondria and normalized to citrate synthase (CS) activity. (A–C) Complex I–IV activities following 1 month of exposure at (A) 0.2 mg/kg BW, (B) 2 mg/kg BW, and (C) 10 mg/kg BW. (D) Complex I–IV activities following 5 months of exposure at 0.2 mg/kg BW. Data are presented as mean ± SEM. Individual symbols represent independent biological replicates. Statistical significance was determined using one-way ANOVA followed by Dunnett’s multiple-comparisons test. *p < 0.05, **p < 0.01, ***p < 0.001, ****p < 0.0001; ns, not significant.

In contrast to the limited changes observed after 1 month at 0.2 mg/kg BW, exposure at the same administered dose for 5 months was associated with significant inhibition of multiple respiratory complexes. GenX significantly reduced all four complexes (control, Complex I–IV: 1.000 ± 0.286, 1.000 ± 0.352, 1.000 ± 0.240, and 1.000 ± 0.244; GenX, Complex I: 0.427 ± 0.055, p = 0.0162; Complex II: 0.358 ± 0.062, p = 0.0063; Complex III: 0.430 ± 0.051, p = 0.0167; Complex IV: 0.457 ± 0.067, p = 0.0239), whereas PFOS significantly reduced only Complex II (0.434 ± 0.032, p = 0.0178), and P+G significantly reduced Complex I, II, and IV (0.486 ± 0.029, p = 0.0242; 0.451 ± 0.015, p = 0.0149; and 0.536 ± 0.016, p = 0.0464, respectively) (Fig. 3D). These findings suggest that the duration of PFAS exposure is important when assessing mitochondrial respiratory enzyme activity. However, direct statistical comparisons between the 1-month and 5-month groups would be required to establish a treatment-duration effect.

#### 3.2.1. Sub-chronic and chronic PFOS and GenX exposure produces different mitochondrial responses in the brain

Mitochondrial Complex I–IV enzyme activities were also measured in isolated brain mitochondria and normalized to citrate synthase activity (Table S2).

Unlike the liver, brain mitochondrial Complex I–IV activities showed no statistically significant changes following 1 month of exposure at any of the three doses tested (Fig. 4A–C). This difference suggests that the mitochondrial responses to repeated PFAS exposure vary between liver and brain under the experimental conditions used.

**Figure 4.**
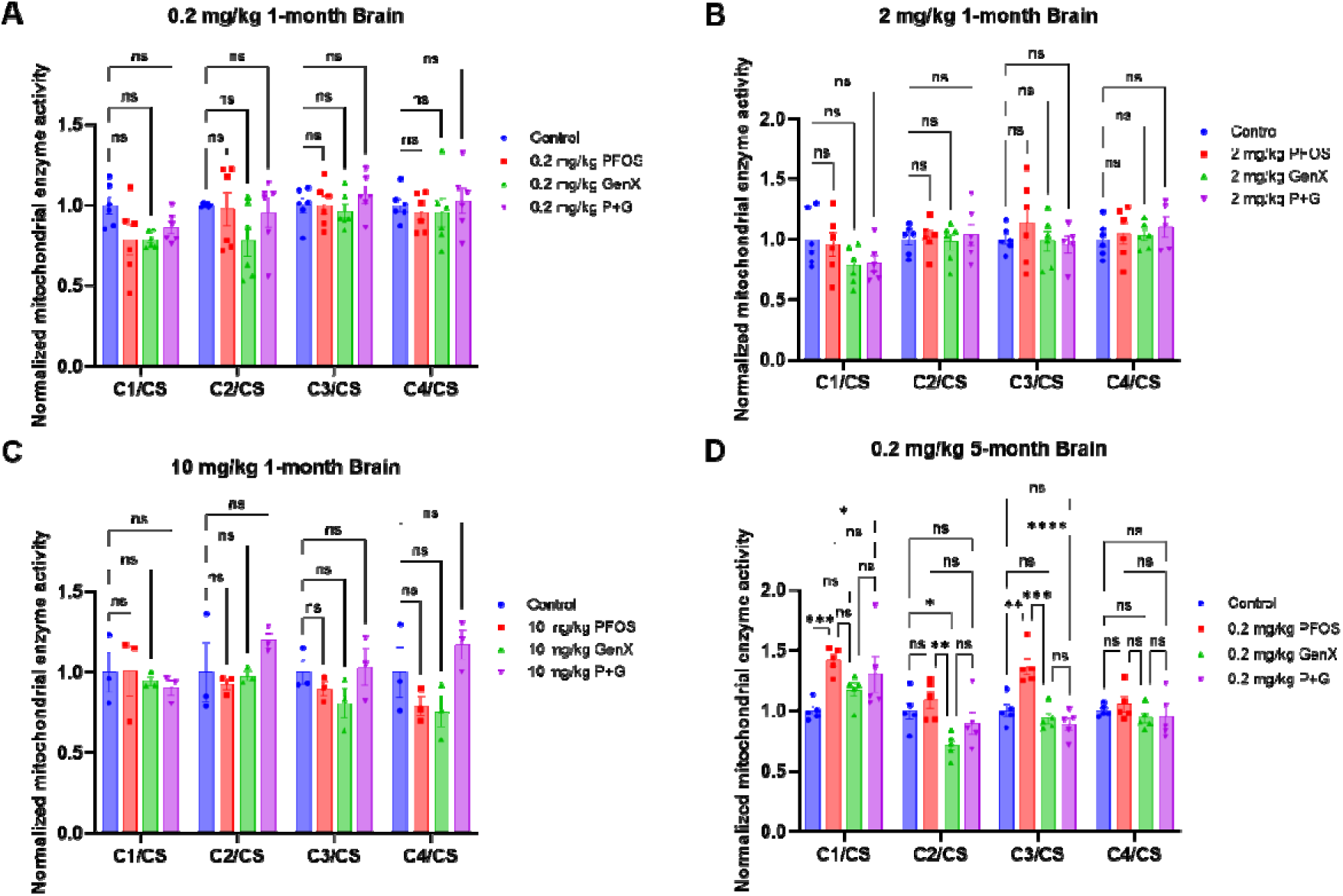
Effects of sub-chronic and chronic PFOS and GenX exposure on mitochondrial respiratory complex activities in mouse brain mitochondria. Mice were exposed to PFOS, GenX, or their combination (P+G) at 0.2, 2, and 10 mg/kg BW for 1 month or at 0.2 mg/kg BW for 5 months. Mitochondrial Complex I–IV activities were measured in isolated brain mitochondria and normalized to citrate synthase (CS) activity. (A–C) Complex I–IV activities following 1 month of exposure at (A) 0.2 mg/kg BW, (B) 2 mg/kg BW, and (C) 10 mg/kg BW. (D) Complex I–IV activities following 5 months of exposure at 0.2 mg/kg BW. Data are presented as mean ± SEM. Individual symbols represent independent biological replicates. Statistical significance was determined using one-way ANOVA followed by Dunnett’s multiple-comparisons test for panels A–C and Tukey’s multiple-comparisons test for panel D. *p < 0.05, **p < 0.01, ***p < 0.001, ****p < 0.0001; ns, not significant.

The 5-month exposure results revealed significant changes in brain Complex I, II, and III activities that were not observed following 1 month of exposure (Fig. 4D). Notably, PFOS increased Complex I and III activities (control, 1.000 ± 0.036 and 1.000 ± 0.053; 1.421 ± 0.048, p = 0.0003 and 1.367 ± 0.070, p = 0.0021, respectively), whereas GenX reduced Complex II activity (control, 1.000 ± 0.071; 0.715 ± 0.046, p = 0.0242). Although repeated exposure produced more extensive changes in liver mitochondrial enzyme activities, the brain showed selective alterations after 5 months.

#### 3.2.2. Combined PFOS and GenX exposure alters body weight following sub-chronic and chronic treatment

Overall, the body weight responses differed across exposure levels and durations. Combined PFOS and GenX exposure produced greater body weight loss than either compound alone following 1 month of treatment at 2 and 10 mg/kg BW (body weight change at 2 mg/kg BW: control, 2.833 ± 0.792 g; P+G, −3.000 ± 0.894 g, p < 0.0001; at 10 mg/kg BW: control, 2.833 ± 0.833 g; PFOS, −5.167 ± 0.654 g, p < 0.0001; GenX, −2.667 ± 0.667 g, p < 0.0001; P+G, −8.000 ± 1.238 g, p < 0.0001), while prolonged exposure at 0.2 mg/kg BW significantly reduced body weight gain (control, 12.200 ± 1.489 g; P+G, 7.500 ± 1.186 g, p = 0.0012). Water intake also showed different responses across exposure conditions, increasing following GenX and P+G exposure at 2 mg/kg BW and decreasing following PFOS and P+G exposure at 10 mg/kg BW. Detailed results are in the Supplementary Information (Section S3, Fig. S1).

### 3.3. PFOS and GenX were detected in plasma and different brain regions following 1 month of exposure

To determine the distribution of PFOS and GenX following repeated exposure, their concentrations were measured in plasma and different brain regions using LC-MS. Mice were treated with PFOS, GenX, or their combination (P+G) at 0.2, 2, and 10 mg/kg BW for 1 month. Brain regions analyzed included the cerebral cortex at all three doses, and the cerebellum and brainstem at 2 mg/kg BW. Detailed values are in the Supplementary Information (Section S4).

The plasma results show that PFOS and GenX concentrations differed between individual and combined treatments depending on exposure level. At 0.2 mg/kg BW, combined exposure was associated with significantly higher plasma PFOS concentrations (8891.667 ± 671.953 ng/mL vs. 4148.333 ± 482.014 ng/mL in the PFOS group, p = 0.0061), whereas at 10 mg/kg BW, plasma GenX concentrations were significantly lower in the P+G group than in the GenX-alone group (3813.000 ± 1774.344 ng/mL vs. 23634.667 ± 666.243 ng/mL, p < 0.0001). Higher co-exposure exhibited a reduction, likely due to the fact that PFOS and GenX both bind to serum albumin [17, 49], and at higher doses the internal levels were competing for binding with serum albumin, resulting in the reduction of the PFAS levels. These results indicate differences in circulating PFAS concentrations between individual and combined exposure under specific experimental conditions (Fig. 5A–F). The lower dose (0.2 mg/kg BW) gave plasma levels about ∼ 2.5X the highest serum level reported in humans [9], and the high dose (2 mg/kg BW) about ∼ 4.5X the highest reported occupational serum level [50].

**Figure 5.**
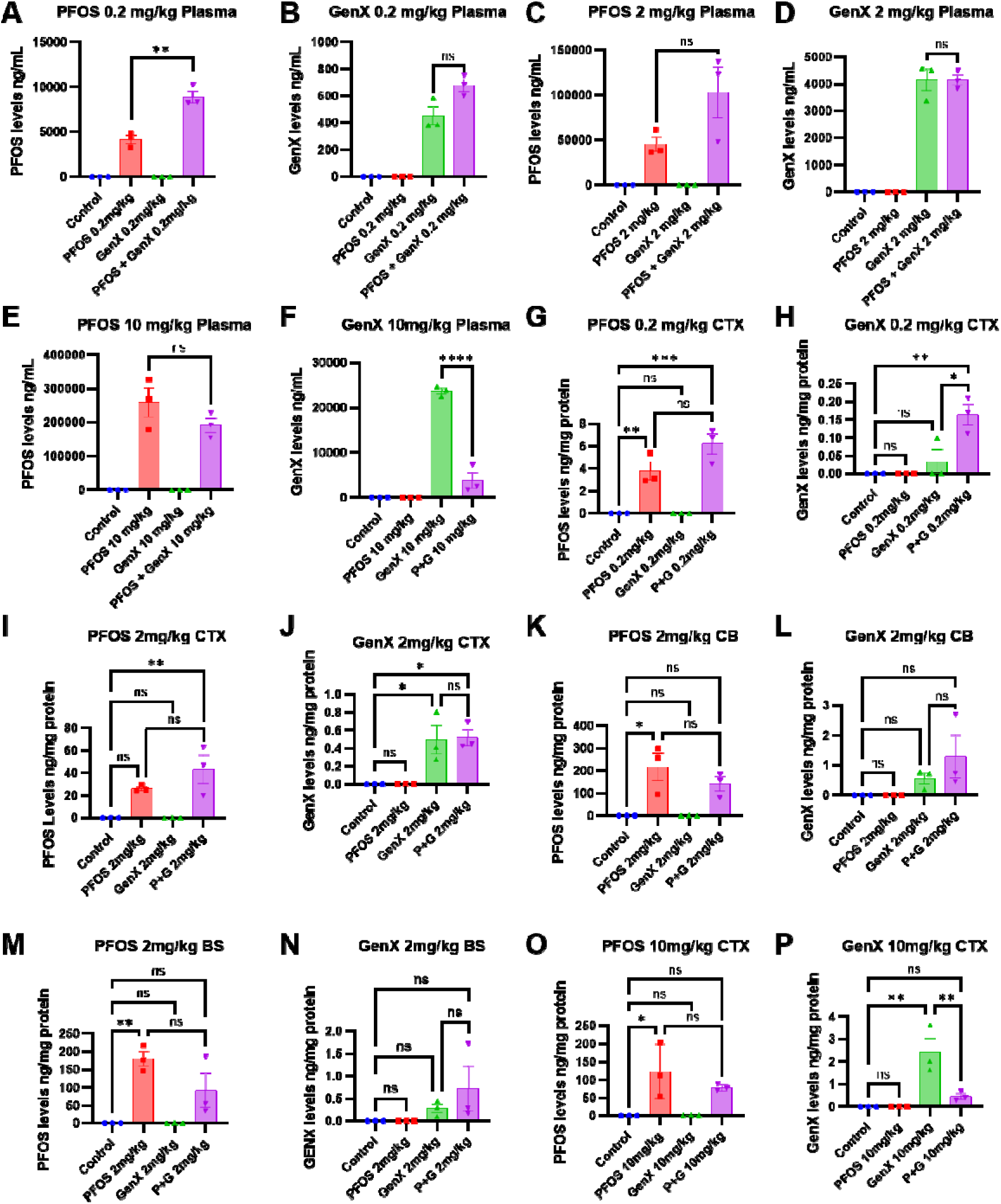
Distribution of PFOS and GenX in mouse plasma and different brain regions following sub-chronic exposure. Mice were exposed to PFOS, GenX, or their combination (P+G) at 0.2, 2, and 10 mg/kg BW for 1 month. PFOS and GenX concentrations were measured using liquid chromatography–mass spectrometry (LC-MS). (A–B) Plasma concentrations of (A) PFOS and (B) GenX following exposure at 0.2 mg/kg BW. (C–D) Plasma concentrations of (C) PFOS and (D) GenX at 2 mg/kg BW. (E–F) Plasma concentrations of (E) PFOS and (F) GenX at 10 mg/kg BW. (G–H) Concentrations of (G) PFOS and (H) GenX in the cerebral cortex (CTX) at 0.2 mg/kg BW. (I–J) Concentrations of (I) PFOS and (J) GenX in the cerebral cortex at 2 mg/kg BW. (K–L) Concentrations of (K) PFOS and (L) GenX in the cerebellum (CB) at 2 mg/kg BW. (M–N) Concentrations of (M) PFOS and (N) GenX in the brainstem (BS) at 2 mg/kg BW. (O–P) Concentrations of (O) PFOS and (P) GenX in the cerebral cortex at 10 mg/kg BW. Plasma concentrations are expressed as ng/mL, whereas brain tissue concentrations are normalized to protein content (ng/mg protein). Data are presented as mean ± SEM. Individual symbols represent independent biological replicates. Statistical significance was determined using one-way ANOVA followed by Tukey’s multiple-comparisons test. *p < 0.05, **p < 0.01, ***p < 0.001, ****p < 0.0001; ns, not significant.

PFOS and GenX were detected in different brain regions following repeated exposure (Fig. 5G–P), indicating that both compounds can reach brain tissue under the experimental conditions used. However, their concentrations varied among brain regions and treatments. At 2 mg/kg BW, average PFOS concentrations were higher in the cerebellum and brainstem than in the cerebral cortex, particularly in the PFOS-alone group. This pattern suggests regional variation in PFOS concentrations, although statistical comparisons between regions are required to establish significant regional differences. GenX was also detected in the cerebral cortex, cerebellum, and brainstem, although its measured concentrations were generally much lower than those of PFOS. At 2 mg/kg BW, GenX uptake in the cerebral cortex was 52 times less than that of PFOS (0.497 ± 0.160 and 25.868 ± 1.695 ng/mg protein, respectively) (Fig. 5I–J). Together, these results demonstrate the presence of both PFOS and GenX in brain tissues following repeated exposure, with differences in measured concentrations across compounds and brain regions.

### 3.4. PFOS and GenX distribution in liver, plasma, and brain regions following subchronic exposure in rats

To examine the tissue distribution of PFOS and GenX following repeated exposure, their concentrations were measured in the liver, plasma, and three brain regions, including the cerebral cortex, cerebellum, and brainstem, using LC-MS. Rats were exposed to PFOS, GenX, or their combination (P+G) at 2 mg/kg BW. Tissue concentrations were compared between individual and combined exposure groups to determine whether co-exposure was associated with differences in PFAS distribution. Detailed values are in the Supplementary Information (Section S5).

Overall, PFOS and GenX showed different distribution patterns across liver, plasma, and brain tissues (Fig. S2). Combined exposure was associated with significantly lower plasma PFOS concentrations compared with PFOS alone (164785.739 ± 13217.041 vs. 198231.007 ± 5064.608 ng/mg protein, p = 0.0411) **[CHECK 10: Rat plasma is reported in ng/mg protein in the Supplementary Information, but values this large look like ng/mL. Confirm the unit.]**, whereas no significant differences between individual and combined treatments were observed in the liver or measured brain regions. These findings suggest that differences in PFAS concentrations between individual and combined exposure may vary across biological compartments.

### 3.5. PFOS and GenX distribution in liver, plasma, and different brain regions

PFOS and GenX were detected in plasma and brain tissue after repeated exposure in both species. Different brain regions were studied in rats (cerebral cortex, cerebellum, and brainstem; Fig. S2), whereas in mice the cerebral cortex was the region analyzed at all three doses, and the cerebellum and brainstem were analyzed only at 2 mg/kg BW (Fig. 5). In mouse plasma, PFOS concentrations were significantly higher in the P+G group than in the PFOS-alone group at 0.2 mg/kg BW (8891.667 ± 671.953 ng/mL vs. 4148.333 ± 482.014 ng/mL, p = 0.0061), and GenX concentrations were significantly lower in the P+G group than in the GenX-alone group at 10 mg/kg BW (3813.000 ± 1774.344 ng/mL vs. 23634.667 ± 666.243 ng/mL, p < 0.0001) (Fig. 5A–F). In the mouse cerebral cortex, GenX concentrations were significantly higher in the P+G group than in the GenX-alone group at 0.2 mg/kg BW (0.163 ± 0.029 ng/mg protein vs. 0.033 ± 0.033 ng/mg protein, p = 0.0135) and significantly lower at 10 mg/kg BW (0.450 ± 0.131 ng/mg protein vs. 2.430 ± 0.615 ng/mg protein, p = 0.0092) (Fig. 5G–H, O–P). At 2 mg/kg BW, PFOS was significantly higher than the control in the cerebellum (217.427 ± 62.712 ng/mg protein, p = 0.0103) and brainstem (179.508 ± 20.539 ng/mg protein, p = 0.0053) of the PFOS group (Fig. 5I–N). In rats, PFOS and GenX were detected in the liver, and co-exposure did not significantly change their concentrations (Fig. S2). Plasma PFOS was significantly lower in the P+G group than in the PFOS-alone group (164785.739 ± 13217.041 vs. 198231.007 ± 5064.608 ng/mg protein, p = 0.0411). PFOS was significantly higher than the control in the cerebral cortex, cerebellum, and brainstem of the PFOS and P+G groups (cerebral cortex, 65.062 ± 16.159, p < 0.0001 and 39.205 ± 2.293, p = 0.0019; cerebellum, 51.853 ± 3.288, p < 0.0001 and 56.074 ± 10.279, p < 0.0001; brainstem, 67.098 ± 6.276, p < 0.0001 and 62.020 ± 10.630, p < 0.0001; ng/mg protein; control, 0.000 ± 0.000), and GenX was significantly higher than the control in the cerebellum and brainstem (cerebellum, 0.214 ± 0.047, p = 0.0221 and 0.315 ± 0.093, p = 0.0006; brainstem, 0.189 ± 0.020, p = 0.0498 and 0.295 ± 0.088, p = 0.0013; ng/mg protein, for GenX and P+G, respectively), with no significant differences between individual and combined exposure in any brain region. Detailed values are in Supplementary Information (Sections S4 and S5).

### 3.6. Sub-chronic PFOS and GenX exposure alters mitochondrial OXPHOS protein expression across different brain regions

To examine the effects of PFOS and GenX exposure on mitochondrial respiratory chain protein expression, we assessed the levels of representative oxidative phosphorylation (OXPHOS) proteins in the cerebral cortex, cerebellum, and brainstem following 1 month of treatment at 2 and 10 mg/kg BW or 5 months of treatment at 0.2 mg/kg BW. The OXPHOS proteins included NADH oxidoreductase subunit B8 (Complex I; NDUFB8), succinate dehydrogenase complex iron-sulfur subunit B (Complex II; SDHB), ubiquinol-cytochrome c reductase core protein 2 (Complex III; UQCRC2), mitochondrially encoded cytochrome c oxidase I (Complex IV; MTCO1), and ATP synthase F1 subunit alpha (Complex V; ATP5F1A). OXPHOS protein expression was assessed by Western blot and normalized to Hsp60. Data are presented as mean ± SEM. Detailed values and Western blot methods are in the Supplementary Information (Section S6).

#### 3.6.1. Cerebral cortex (2 mg/kg BW for 1 month)

At 2 mg/kg BW, Complex I protein expression was significantly increased in the PFOS (1.980 ± 0.263, p < 0.0001) and GenX (1.714 ± 0.115, p = 0.0006) groups compared with the control (1.099 ± 0.146), whereas no significant difference was observed in the P+G group (Fig. 6A–B). Complex II and Complex V expression were significantly increased in the GenX group (1.396 ± 0.094, p = 0.0494 and 1.355 ± 0.135, p = 0.0467, respectively; control, 1.022 ± 0.055 and 1.000 ± 0.053). Complex III expression was significantly reduced in the PFOS group (0.372 ± 0.101, p = 0.0039; control, 0.956 ± 0.044), and Complex IV expression was significantly reduced in the PFOS (0.193 ± 0.124, p = 0.0011) and GenX (0.342 ± 0.102, p = 0.0100) groups (control, 1.000 ± 0.118). No significant changes were observed in the P+G group.

**Figure 6.**
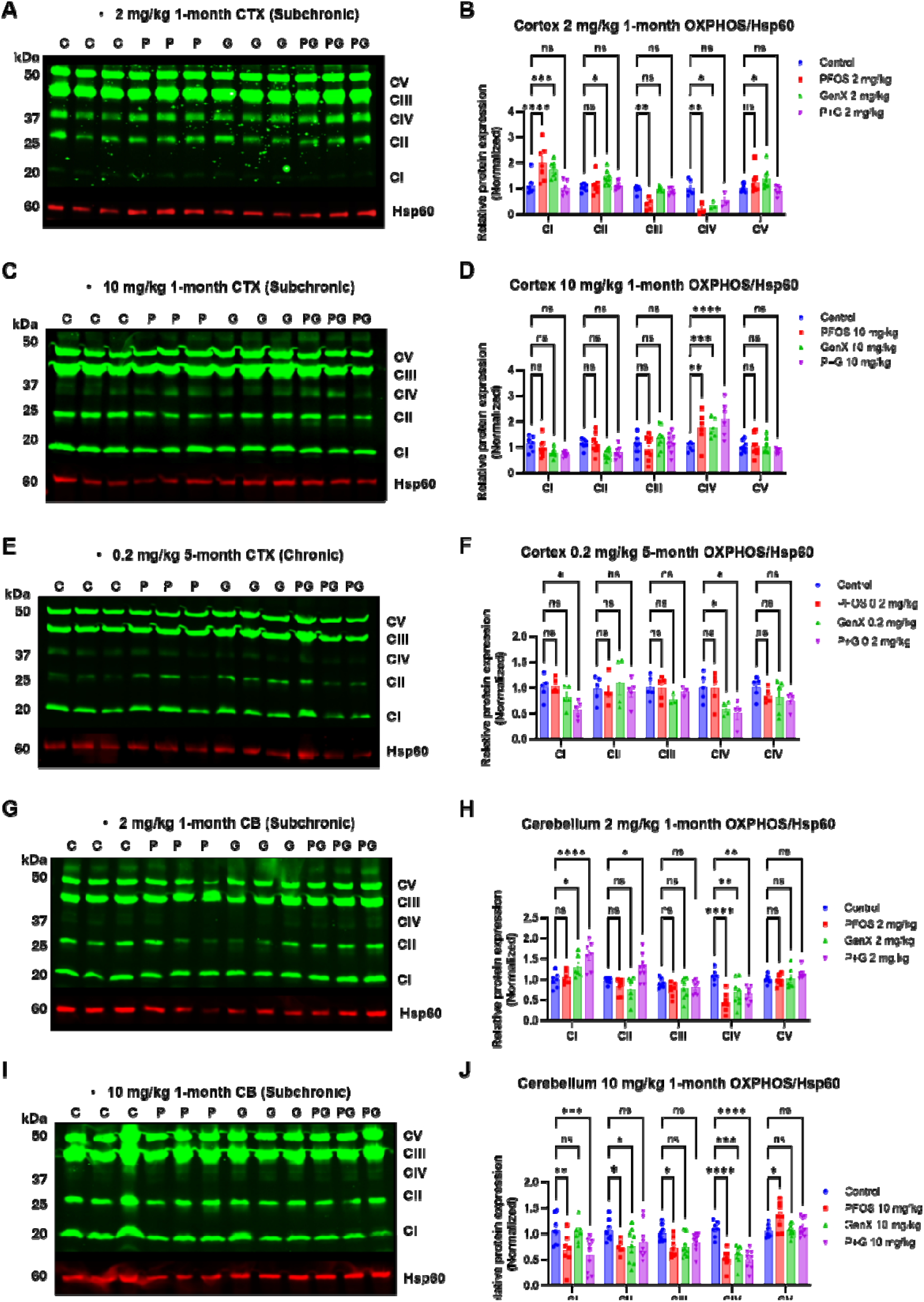
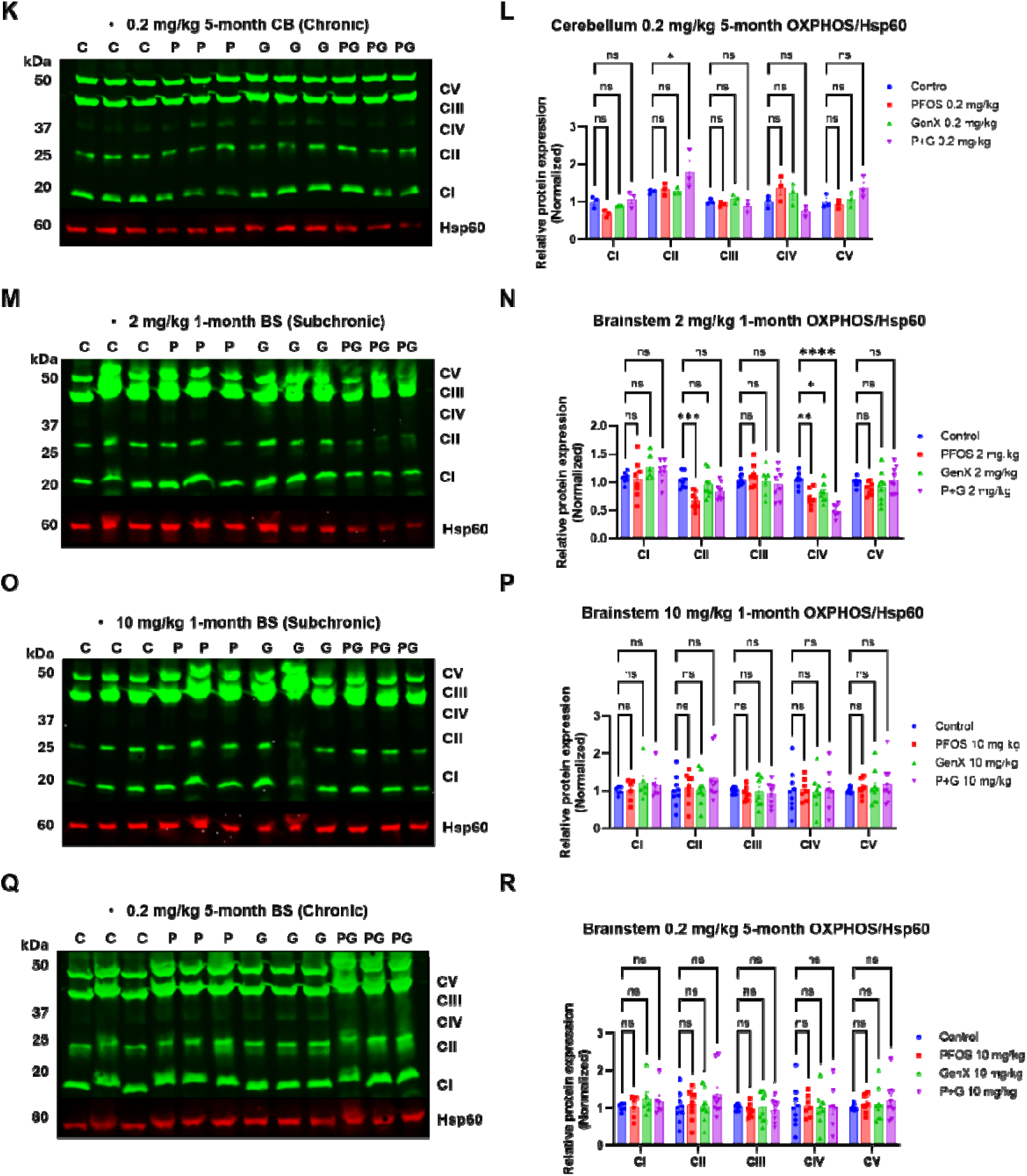
Effects of sub-chronic and chronic PFOS and GenX exposure on mitochondrial oxidative phosphorylation (OXPHOS) protein expression in different mouse brain regions. Mice were exposed to PFOS, GenX, or their combination (P+G) at 2 and 10 mg/kg BW for 1 month or at 0.2 mg/kg BW for 5 months. OXPHOS protein expression was assessed by Western blotting in the cerebral cortex, cerebellum, and brainstem. Representative OXPHOS Western blots and corresponding quantitative analyses are shown for the cerebral cortex following (A–B) 2 mg/kg BW for 1 month, (C–D) 10 mg/kg BW for 1 month, and (E–F) 0.2 mg/kg BW for 5 months; the cerebellum following (G–H) 2 mg/kg BW for 1 month, (I–J) 10 mg/kg BW for 1 month, and (K–L) 0.2 mg/kg BW for 5 months; and the brainstem following (M–N) 2 mg/kg BW for 1 month, (O–P) 10 mg/kg BW for 1 month, and (Q–R) 0.2 mg/kg BW for 5 months. Representative OXPHOS subunits included Complex I (NDUFB8), Complex II (SDHB), Complex III (UQCRC2), Complex IV (MTCO1), and Complex V (ATP5F1A). Protein expression was normalized to Hsp60 and expressed relative to the control group. Data are presented as mean ± SEM. Individual symbols represent independent biological replicates. Statistical significance was determined using two-way ANOVA followed by Dunnett’s multiple-comparisons test. *p < 0.05, **p < 0.01, ***p < 0.001, ****p < 0.0001; ns, not significant.

#### 3.6.2. Cerebral cortex (10 mg/kg BW for 1 month)

At 10 mg/kg BW, Complex IV protein expression was significantly increased in all treatment groups compared with the control (control, 1.000 ± 0.069; PFOS, 1.724 ± 0.263, p = 0.0011; GenX, 1.735 ± 0.153, p = 0.0009; P+G, 2.087 ± 0.280, p < 0.0001) (Fig. 6C–D). No statistically significant differences were observed in Complex I, II, III, or V expression.

#### 3.6.3. Cerebral cortex (0.2 mg/kg BW for 5 months)

To examine the effects of longer-term PFAS exposure, OXPHOS protein expression was also assessed following treatment at 0.2 mg/kg BW for 5 months (Fig. 6E–F).

Following 5 months of exposure at 0.2 mg/kg BW, combined PFOS and GenX exposure significantly reduced Complex I (0.562 ± 0.078, p = 0.0182; control, 1.000 ± 0.105) and Complex IV (0.505 ± 0.133, p = 0.0112; control, 1.000 ± 0.122) protein expression, whereas GenX alone significantly reduced Complex IV expression (0.587 ± 0.070, p = 0.0415). In contrast, PFOS alone did not significantly alter any of the five OXPHOS proteins examined. These results show that chronic exposure at a lower dose was associated with selective changes in mitochondrial OXPHOS protein abundance, particularly following GenX and combined exposure. Across the three exposure conditions, the cerebral cortex showed distinct patterns of OXPHOS protein expression. At 2 mg/kg BW for 1 month, several complexes were affected by PFOS and GenX individually, whereas at 10 mg/kg BW, the significant changes were restricted to increased Complex IV expression. After 5 months at 0.2 mg/kg BW, reduced Complex I and IV expression were observed following combined exposure. These findings suggest that mitochondrial OXPHOS protein expression varies with the compound, exposure level, and duration.

#### 3.6.4. Cerebellum (2 mg/kg BW for 1 month)

At 2 mg/kg BW, Complex I protein expression was significantly increased in the GenX (1.296 ± 0.116, p = 0.0269) and P+G (1.579 ± 0.131, p < 0.0001) groups compared with the control (0.972 ± 0.072), whereas PFOS did not significantly alter Complex I expression (Fig. 6G–H). Complex II expression was significantly increased in the P+G group (1.278 ± 0.107, p = 0.0127; control, 0.966 ± 0.027). Complex IV expression was significantly reduced in all treatment groups (control, 1.081 ± 0.066; PFOS, 0.436 ± 0.080, p < 0.0001; GenX, 0.670 ± 0.115, p = 0.0024; P+G, 0.627 ± 0.075, p = 0.0010). No significant differences were observed in Complex III or V expression.

These results show that PFOS, GenX, and P+G altered OXPHOS protein expression in the cerebellum after 1 month of exposure. Complex IV was significantly reduced following all three treatments, whereas Complex I was increased in the GenX and P+G groups. Combined exposure also increased Complex II expression, which was not significantly altered by either compound alone. Notably, the significant reduction in Complex IV expression was observed following both individual and combined treatments, while the other complexes showed treatment-specific changes.

#### 3.6.5. Cerebellum (10 mg/kg BW for 1 month)

At 10 mg/kg BW, PFOS exposure significantly reduced Complex I–IV protein expression (control, Complex I–V: 1.057 ± 0.100, 1.057 ± 0.075, 1.000 ± 0.051, 1.083 ± 0.059, and 1.053 ± 0.042; PFOS, Complex I: 0.688 ± 0.127, p = 0.0064; Complex II: 0.740 ± 0.056, p = 0.0310; Complex III: 0.729 ± 0.065, p = 0.0430; Complex IV: 0.515 ± 0.063, p < 0.0001) while increasing Complex V expression (1.325 ± 0.093, p = 0.0497) (Fig. 6I–J). GenX significantly reduced Complex II and IV expression (Complex II: 0.759 ± 0.111, p = 0.0226; Complex IV: 0.595 ± 0.067, p = 0.0002), whereas combined exposure significantly reduced Complex I and IV expression (Complex I: 0.581 ± 0.113, p = 0.0001; Complex IV: 0.509 ± 0.071, p < 0.0001). Compared with the 2 mg/kg BW groups, where Complex I expression increased following GenX and P+G exposure, significant reductions in Complex I expression were observed in the PFOS and P+G groups at 10 mg/kg BW. Although the responses differed among treatments, Complex IV expression was significantly reduced in all treatment groups at both exposure levels.

#### 3.6.6. Cerebellum (0.2 mg/kg BW for 5 months)

Following 5 months of exposure at 0.2 mg/kg BW, combined PFOS and GenX exposure significantly increased Complex II protein expression (1.794 ± 0.307, p = 0.0106; control, 1.268 ± 0.034; Fig. 6K–L), whereas neither compound alone produced significant changes in any of the five OXPHOS proteins examined. Although Complex IV expression was numerically higher in the PFOS and GenX groups and lower in the P+G group, these differences were not statistically significant.

Compared with the 1-month exposure groups, where Complex IV expression was significantly reduced at both 2 and 10 mg/kg BW, no significant change in Complex IV expression was observed after 5 months at 0.2 mg/kg BW. Because both dose and exposure duration differed, these observations describe differences across experimental conditions rather than an established duration-dependent response.

#### 3.6.7. Brainstem (2 mg/kg BW for 1 month)

At 2 mg/kg BW, Complex II protein expression was significantly reduced in the PFOS group (p = 0.0004), and Complex IV expression was significantly reduced in all treatment groups compared with the control (PFOS, p = 0.0041; GenX, p = 0.0445; P+G, p < 0.0001) (Fig. 6M–N). No significant differences were observed in Complex I, III, or V expression.

These results show that Complex IV protein expression was consistently reduced following individual and combined PFAS exposure, whereas Complex II was significantly reduced only in the PFOS group.

#### 3.6.8. Brainstem (10 mg/kg BW for 1 month)

At 10 mg/kg BW, no statistically significant differences were observed in Complex I–V protein expression following PFOS, GenX, or P+G exposure (Fig. 6O-P).

In contrast with the 2 mg/kg BW groups, where Complex II and IV expression were significantly reduced in selected treatment groups, no significant changes in OXPHOS protein expression were detected at 10 mg/kg BW.

#### 3.6.9. Brainstem (0.2 mg/kg BW for 5 months)

Following 5 months of exposure at 0.2 mg/kg BW, combined PFOS and GenX exposure significantly increased Complex IV protein expression (1.697 ± 0.257, p = 0.0022; control, 1.000 ± 0.220; Fig. 6Q–R), whereas neither compound alone produced significant changes in any of the five OXPHOS subunits examined. The brainstem showed selective alterations in OXPHOS protein expression following PFOS and GenX exposure. At 2 mg/kg BW for 1 month, Complex IV expression was reduced following individual and combined treatments, while Complex II expression was reduced only following PFOS exposure. At 10 mg/kg BW, no significant changes were detected in any of the five OXPHOS proteins. In contrast, chronic exposure at 0.2 mg/kg BW was associated with increased Complex IV expression following combined treatment.

### 3.7. Molecular docking analysis reveals different predicted interactions of PFOS and GenX with mitochondrial Complexes III and IV

Molecular docking was performed to investigate the potential interactions of PFOS and GenX, individually and in combination, with mitochondrial Complexes III and IV using human and mouse protein structures. The human docking configurations are shown in Fig. 7. Docking calculations were completed for all 18 species–complex–site–ligand conditions. The predicted docking scores are presented in Table S3.

**Figure 7.**
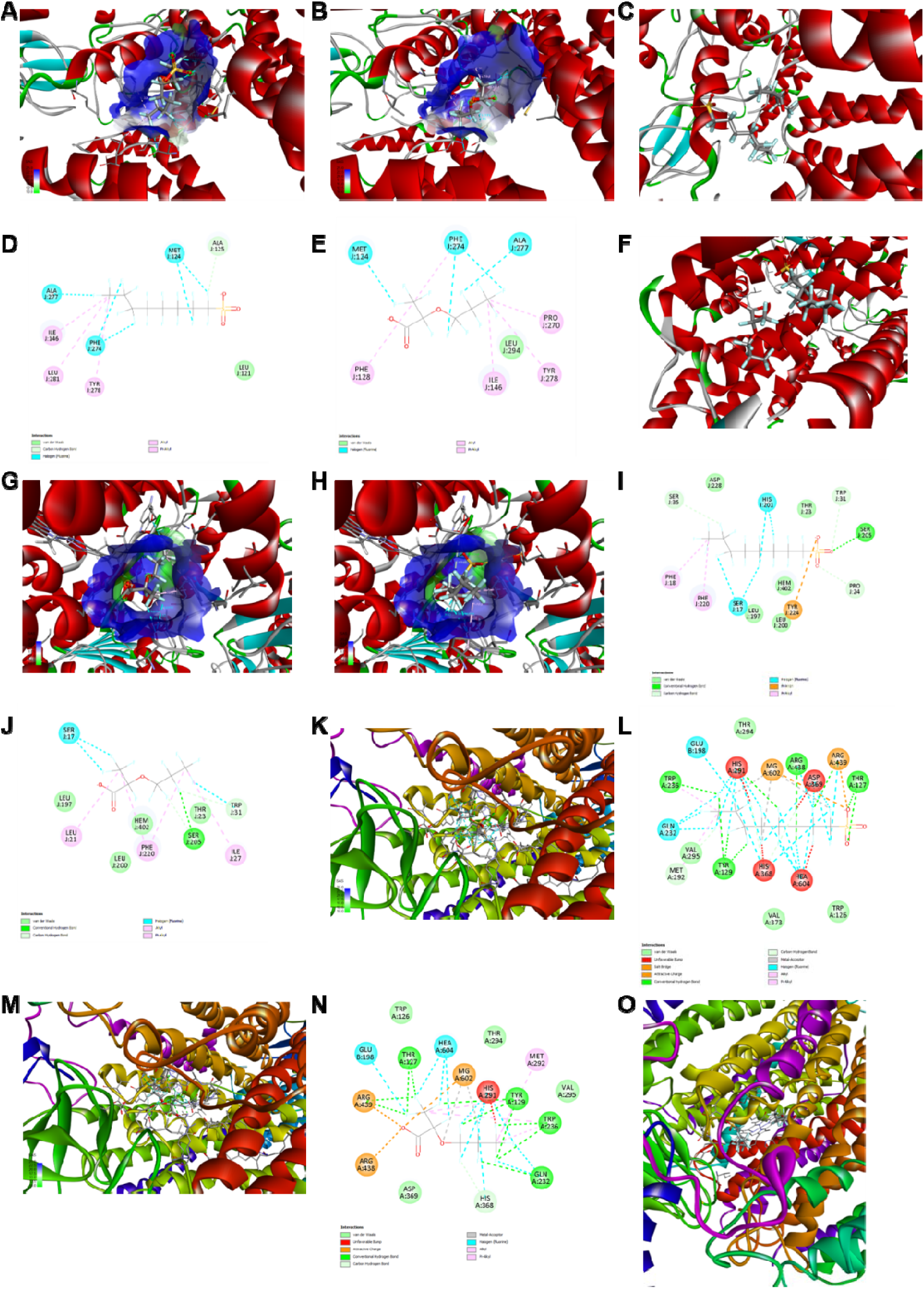
Molecular docking configurations and predicted protein-ligand interactions of PFOS and GenX with human mitochondrial Complex III and IV. Molecular docking was performed at the Qo and Qi binding sites of human mitochondrial Complex III (PDB ID: 9HZL; A-J) and at the catalytic region of human mitochondrial Complex IV (PDB ID: 5Z62; K-O). (A-E) Qo binding site: (A) PFOS docking pose, (B) GenX docking pose, (C) simultaneous PFOS-GenX docking configuration, (D) PFOS 2D interaction diagram, and (E) GenX 2D interaction diagram. (F-J) Qi binding site: (F) PFOS docking pose, (G) GenX docking pose, (H) simultaneous PFOS-GenX docking configuration, (I) PFOS 2D interaction diagram, and (J) GenX 2D interaction diagram. (K-O) Human Complex IV: (K) three-dimensional docking pose of PFOS, (L) corresponding two-dimensional protein–ligand interaction diagram of PFOS, (M) three-dimensional docking pose of GenX, (N) corresponding two-dimensional protein-ligand interaction diagram of GenX, and (O) simultaneous PFOS-GenX docking configuration. Three-dimensional structures were visualized using BIOVIA Discovery Studio Visualizer, with protein secondary structures represented as ribbons and ligands and surrounding amino acid residues displayed as sticks. Two-dimensional diagrams illustrate predicted interactions with neighboring amino acid residues, including hydrophobic, van der Waals, and software-assigned hydrogen-bond and halogen-associated contacts.

For Complex III (Fig. 7A-J), PFOS generally exhibited more favorable predicted docking scores than GenX, except at the mouse Qi site, where GenX scored slightly more favorably. In human Complex III, both compounds exhibited more favorable scores at Qi than Qo. Simultaneous PFOS-GenX docking also generated joint configurations at all four Complex III binding regions. The strongest individual PFOS score occurred at the human Qi site (−8.711 kcal/mol), while the strongest individual GenX score also occurred at human Qi (−7.407 kcal/mol). The mouse Qi site was an exception to the general PFOS-GenX pattern, with GenX scoring slightly more favorably than PFOS

In contrast, Complex IV produced substantially different docking scores (Fig. 7K-O). At the human heme a₃-CuB region, PFOS exhibited a positive docking score of +5.146 kcal/mol, whereas GenX showed a negative score of −1.914 kcal/mol. Similarly, in mouse Complex IV, GenX exhibited a more favorable docking score (−5.160 kcal/mol) than PFOS (−0.1504 kcal/mol). These findings indicate that GenX received more favorable predicted docking scores than PFOS at the selected Complex IV catalytic regions in both species. The positive human PFOS score suggests an unfavorable predicted docking configuration under the selected docking conditions

These findings indicate differences in predicted docking scores between respiratory complexes, species, and the individual PFAS compounds.

For simultaneous PFOS-GenX docking, joint configurations were obtained at all four Complex III binding regions and both Complex IV docking regions. The predicted scores ranged from −12.070 kcal/mol at the human Complex III Qi site to +9.454 kcal/mol at the human Complex IV heme a₃–CuB region. Because each mixture calculation contains two ligands, the scores cannot be directly compared with individual ligand scores to establish synergistic or antagonistic binding.

## 4. Discussion

The present study demonstrated that the effects of PFOS and GenX on mitochondrial respiratory complex activities vary depending on exposure levels, duration, and tissue. Acute exposure to PFOS and GenX inhibited mitochondrial Complex III and IV activities in isolated liver and brain mitochondria, with combined exposure producing greater inhibition than either compound alone under several experimental conditions. However, chronic and sub-chronic exposure resulted in different mitochondrial responses, particularly between the liver and brain. While higher exposure concentrations were associated with extensive inhibition of liver mitochondrial complexes, prolonged exposure at a lower concentration produced selective changes in mitochondrial enzyme activities. These findings suggest that mitochondrial responses to PFOS and GenX are influenced by both exposure conditions and tissue-specific biological processes.

The inhibition of mitochondrial Complex III and IV activities following acute PFOS and GenX exposure suggests that these compounds alter the activities of enzymes involved in the electron transport chain. Previous studies from our group demonstrated that PFOS inhibited mitochondrial respiratory complexes II–IV in isolated rat liver mitochondria [30]. Similarly, our previous research showed that GenX significantly reduced Complex IV activity at 75 ppm in isolated rat brain mitochondria [31]. In the present study, PFOS also inhibited Complex III and IV activities in isolated mouse liver and brain mitochondria, whereas GenX exhibited different concentration-dependent responses. Specifically, PFOS significantly inhibited Complex IV activity at all concentrations tested in isolated mouse liver mitochondria, whereas GenX produced significant inhibition only at 75 ppm. Furthermore, combined exposure produced greater inhibition than either compound alone at several concentrations.

Complex III and IV are essential for electron transport and the maintenance of the proton gradient required for ATP synthesis [23, 24]. Disruption of these enzymes may therefore interfere with mitochondrial energy metabolism. A recent study reported that PFOS exposure reduced mitochondrial oxidative phosphorylation-related gene expression in human liver cells, including genes encoding NADH dehydrogenase, cytochrome c oxidase, and ATP synthase subunits. These transcriptional changes were accompanied by increased ROS, reduced ATP levels, and apoptosis [38]. The present study measured respiratory enzyme activities and OXPHOS protein levels. These findings provide additional evidence that PFOS can affect multiple components of mitochondrial energy metabolism.

The mitochondrial responses observed following repeated exposure were more complex than those following acute treatment. At 0.2 mg/kg BW, combined PFOS and GenX exposure significantly increased liver Complex I activity after 1 month, whereas higher exposure concentrations produced more extensive inhibition of respiratory complexes. The increased Complex I activity at the lowest concentration may reflect changes in mitochondrial enzyme regulation during repeated exposure. However, increased enzyme activity does not necessarily indicate enhanced oxidative phosphorylation, and whether this response represents a compensatory mechanism remains unclear. In contrast, at 2 and 10 mg/kg BW, several complexes showed reduced activities, suggesting that higher PFAS exposure concentrations are associated with broader disturbances in mitochondrial respiratory enzyme function.

One particularly interesting finding was that GenX significantly reduced all four mitochondrial respiratory complex activities in the liver at 2 mg/kg BW, whereas PFOS alone significantly inhibited only Complex III. This difference suggests that GenX can affect hepatic mitochondrial function despite its relatively rapid elimination compared with PFOS and that the effect of GenX and PFOS are specific, not generic. At 10 mg/kg BW, PFOS, GenX, and their combination significantly reduced Complex I–IV activities in liver mitochondria. The broader inhibition observed at this concentration may reflect more extensive mitochondrial disturbances compared with those observed at lower exposure concentrations. A recent study reported that PFOS exposure altered mitochondrial respiratory function and metabolic pathways in mice, although the affected pathways differed between liver and skeletal muscle. PFOS reduced Complex II-linked respiratory capacity in the liver, whereas Complex I- and IV-associated respiratory capacity was affected in skeletal muscle [39]. These findings suggest that PFOS can affect mitochondrial energy metabolism through different pathways depending on the tissue examined. However, the reductions in Complex I–IV activities observed in the present study do not establish whether these effects result from direct inhibition of respiratory enzymes, changes in mitochondrial protein expression, or indirect effects on mitochondrial function.

The differences in mitochondrial responses between the liver and brain may reflect differences in tissue-specific metabolic regulation and internal PFAS exposure. A previous study reported that PFOS induced different transcriptomic and mitochondrial responses in the liver and skeletal muscle, despite higher PFOS accumulation in the liver [39]. Although that study did not examine the brain, its findings suggest that mitochondrial responses to PFOS may depend on tissue-specific biological characteristics rather than tissue concentrations alone. Furthermore, a recent study using human induced pluripotent stem cell (iPSC)-derived cortical neurons demonstrated different cellular responses to PFOA and GenX exposure [40]. PFOA exposure caused persistent mitochondrial dysfunction, whereas GenX exposure was associated with long-lasting alterations in neuronal network morphology and increased susceptibility to autolysosomal stress. Although this study examined PFOA rather than PFOS, the findings suggest that different PFAS may affect neuronal function through different cellular mechanisms. Therefore, the absence of significant changes in brain mitochondrial complex activities following shorter exposure does not necessarily exclude alterations in other aspects of mitochondrial or neuronal function.

Differences in internal PFAS exposure may partly explain the greater mitochondrial effects in the liver than in the brain. In rodents, PFOS accumulates mainly in the liver, with far more in the liver than in serum, whereas only moderate quantities are found in the brain[41]. GenX shows a similar pattern, with the highest concentrations in the liver and about 40 times less in the brain after a single oral dose[42]. Although PFOS does cross the blood–brain barrier and brain concentrations increase with dose, accumulation also differs among brain regions[43]. This is consistent with our observation that PFOS concentrations varied among the cerebral cortex, cerebellum, and brainstem. However, tissue dose alone does not fully account for our findings. GenX concentrations in the cerebral cortex were much lower than those of PFOS, yet GenX still altered OXPHOS protein expression in the same region (Fig. 6A–B).

Furthermore, liver and brain also differ in how their mitochondria function. Hepatic mitochondria carry out fatty acid β-oxidation, ATP production, and redox maintenance, and PFAS are known to disturb these processes in the liver[44]. PPARα, which increases fatty acid oxidation and may lead to oxidative stress and lipid accumulation in hepatocytes, is activated by both PFOS[45, 46] and GenX[47, 48]. In contrast, the brain relies mainly on glucose, and its capacity for fatty acid oxidation is relatively limited[49]. Hence, the liver may therefore be exposed to an additional metabolic route of toxicity uncommon in the brain. Yet, it remains unclear whether PFOS inhibits mitochondrial β-oxidation or acts mainly through PPARα. One study found that PFOA and PFOS did not change palmitate-supported respiration in isolated mitochondria or human hepatocytes[12]. Our data do not distinguish between these mechanisms, and the direct effects observed in isolated liver and brain mitochondria after acute exposure suggest that tissue dose and PPARα signaling are not the only factors involved.

Additionally, cellular composition of the brain may also contribute to the limited changes in brain mitochondrial complex activities. Brain mitochondria isolated from whole tissue come from neurons, astrocytes, and microglia, which differ in their energy metabolism and requirements. While neurons depend mainly on oxidative phosphorylation and have low glycolytic capacity, astrocytes rely more on glycolysis[50]. Complex I is largely assembled into super-complexes in neurons, whereas astrocytes contain a higher fraction of free Complex I and produce more ROS[51]. Furthermore, microglia depend on oxidative phosphorylation under resting conditions and shift to glycolysis upon inflammatory activation[52]. Because isolated brain mitochondria represent a mixture mitochondria extracted from these three cell types, a change in one population may be diluted or masked in measurements of total enzyme activity. This may be relevant for PFOS, which has been reported to disrupt the microglial transcriptome and increase inflammation in microglia[53]. However, we did not examine cell-type-specific mitochondrial responses, and these possibilities will need to be tested in future studies, for example in isolated neurons, astrocytes, and microglia exposed to PFOS and GenX.

The increased brain Complex I and III activities following chronic PFOS exposure may reflect changes in mitochondrial enzyme regulation. However, these increases do not necessarily indicate improved mitochondrial function, as enzyme activity measurements alone cannot determine whether they represent adaptive responses or mitochondrial dysregulation. Further assessment of mitochondrial respiration, ATP production, and oxidative stress is needed to clarify their functional significance.

The analysis of PFOS and GenX distribution provided further information about differences in internal levels. Both compounds were detected in plasma and brain tissues following repeated exposure, with PFOS generally exhibiting higher concentrations than GenX. PFOS concentrations also varied among the cerebral cortex, cerebellum, and brainstem, suggesting differential affinity in brain regions. Previous studies have reported differences in PFAS accumulation across human brain regions and in brain-to-plasma concentration ratios [16, 17]. However, direct statistical comparisons are needed to confirm regional differences, and measurements at a single time point cannot determine PFAS uptake or elimination kinetics. Moreover, the detection of PFOS and GenX in brain tissues does not necessarily indicate corresponding changes in mitochondrial function, as cellular susceptibility and intracellular distribution may influence biological responses.

Binding of both PFOS [17] and GenX [49] to serum albumin may result in competition for albumin binding at the higher dose. Notably, GenX uptake in the cerebral cortex at 2 mg/kg BW was 52 times less than that of PFOS (Fig. 5I–J), yet it was able to alter OXPHOS protein expression in the same region (Fig. 6A–B). This finding is significant, since GenX is perceived to be less toxic. The longer-duration, low-dose exposure is closer to what humans might be exposed to. Given the internal levels observed, a higher dose is necessary because elimination rates differ starkly between rodents and humans [4, 18].

In addition to mitochondrial alterations, combined PFOS and GenX exposure produced greater reductions in body weight than either compound alone at 2 and 10 mg/kg BW following 1 month of exposure. Prolonged combined exposure at 0.2 mg/kg BW also reduced body weight gain, whereas individual treatments did not significantly affect this endpoint. These findings suggest that combined exposure may have greater effects on body weight than individual compounds. Corroborating our findings, previous research has also reported PFOS-associated reductions in body weight and alterations in energy metabolism in mice [39]. However, changes in body weight cannot be directly linked to mitochondrial respiratory complex inhibition, as combined exposure did not consistently produce the greatest reductions in enzyme activities. Differences in water consumption, PFAS intake, and systemic metabolism may also contribute to these observations.

Combined PFOS and GenX exposure produced different mitochondrial responses depending on the exposure conditions. During acute exposure, combined treatment generally caused greater inhibition of Complex III and IV activities than either compound alone. However, following repeated exposure, GenX alone produced comparable or greater reductions in several hepatic respiratory enzyme activities. These findings suggest that the effects of combined exposure may vary depending on exposure duration, tissue, and mitochondrial enzymes. Previous studies have reported PFAS mixture toxicity consistent with concentration addition at low effect levels [33], as well as alterations in mitochondrial function and metabolic pathways following mixture exposure [34]; however, the present findings do not establish additive or synergistic interactions between PFOS and GenX. Although this study provides insights into the effects of PFOS and GenX exposure on mitochondria. Still some caveats should be acknowledged. Measurements of complex mitochondrial activities alone cannot fully characterize mitochondrial function, and differences between exposure durations require further investigation. Future studies incorporating mitochondrial respiration, oxidative stress, and metabolic profiling would help clarify the mechanisms underlying PFOS- and GenX-induced mitochondrial responses observed in this study.

To further explore potential interactions with the mitochondrial respiratory complexes, molecular docking was conducted. The analysis revealed distinct predicted interaction patterns for PFOS and GenX across mitochondrial Complexes III and IV. PFOS generally exhibited more favorable docking scores at Complex III binding sites, whereas GenX showed more favorable scores at the selected Complex IV catalytic regions. Notably, PFOS is a relatively larger molecule than GenX, and dynamics tend to vary when running comparisons. Future *in silico* studies will aim for molecular dynamics simulations to get a better picture, along with the specific inhibitors for the mitochondrial enzymes.

## 5. Conclusion

This study examined the individual and combined effects of PFOS and GenX under acute, sub-chronic, and chronic exposure conditions, focusing on mitochondrial complex activities in the liver and brain. Our findings demonstrate that PFOS and GenX can alter mitochondrial respiratory complex activities, with responses varying according to the compound, tissues and exposure conditions ranging from exposure levels to co-exposures. One particularly important aspect of this study was the effect of co-exposure, which is likely to happen in real-life scenarios. The assessment of PFOS distribution across different brain regions also revealed differences in regional brain accumulation and is especially important with respect to plausible specific disease pathology depending on the brain regions responsible for neurodegenerative disorders. Molecular docking, though preliminary, identified differences in predicted interactions of PFOS and GenX with mitochondrial Complexes III and IV, identifying compound-specific interactions that need to be further tested through molecular dynamic simulations. These findings highlight the importance of considering exposure duration, tissue distribution, and combined PFAS exposure when evaluating their potential toxicological effects. These studies will pave way for in-depth mechanistic studies evaluating PFOS- and GenX-induced mitochondrial alterations and the long-term consequences of exposure to their mixtures.

## Author contribution

SRS conceived the study and performed the Western blot experiments. NK and NB handled the animals for the first experimental batch, and DK and NB handled the animals for the second experimental batch. SP performed all *in silico* studies. JOK performed sample preparation, extraction, and LC–MS/MS analysis for PFAS quantification. NB led the preparation of the original manuscript draft, with DK contributing to the writing in a supporting role. SRS edited the MS and serves as the corresponding author.

## Notes

### Competing Interest Statement

The authors have declared no competing interest.

### Summary of Updates

The MS has been refined and some of the information has been move to supplementary information

