## Supplementary information for "PFOS and GenX Induce Mitochondrial Toxicity and Differentially Alter Respiratory Complex Activities in Liver and Brain"

**S1. Acute ex vivo exposure: detailed Results (liver and brain mitochondria, Complex III and IV)**

PFOS significantly reduced Complex III activity at all concentrations tested, with normalized activity decreasing to 0.7887 ± 0.03601 at 25 ppm (*p* = 0.0205), 0.6769 ± 0.05361 at 50 ppm (***p* = 0.0007), and 0.6067 ± 0.05196 at 75 ppm (****p* < 0.0001), compared with the control (1.000 ± 7.071 × 10⁻⁶) (Fig. 1A). GenX also reduced Complex III activity, although the reduction was not statistically significant at 25 ppm (0.8564 ± 0.05433, *p* > 0.05). Significant reductions were observed at 50 ppm (0.7977 ± 0.01661, **p* = 0.0234) and 75 ppm (0.6068 ± 0.02173, ****p* = 0.0002) compared with the control (Fig. 1B).

Combined exposure to PFOS and GenX resulted in lower mean Complex III activity compared with either compound alone at the concentrations tested. Complex III activity decreased from 1.000 ± 5.000 × 10⁻⁶ in the control group to 0.6534 ± 0.06563 at 25 ppm (****p* = 0.0007), 0.3695 ± 0.05804 at 50 ppm (*****p* < 0.0001), and 0.3142 ± 0.06653 at 75 ppm (*****p* < 0.0001) (Fig. 1C).

To compare the effects of individual and combined treatments at the same concentration, Complex III activity was further analyzed at 25, 50, and 75 ppm (Fig. 1D–F). At 25 ppm, PFOS significantly reduced Complex III activity to 0.7887 ± 0.03601 (**p* = 0.0194), whereas GenX showed a reduction that was not statistically significant (0.8564 ± 0.05433, *p* = 0.1468), compared with the control (1.000 ± 5.000 × 10⁻⁶). The P+G group also showed a significant reduction in Complex III activity (0.6534 ± 0.06563, ****p* = 0.0006) compared with the control. However, no statistically significant differences were detected among the PFOS, GenX, and P+G treatment groups at this concentration (Fig. 1D).

At 50 ppm, Complex III activity was significantly reduced by PFOS (0.6769 ± 0.05361, ****p* = 0.0005), GenX (0.7977 ± 0.01661, **p* = 0.0204), and P+G (0.3695 ± 0.05804, *****p* < 0.0001), compared with the control (1.000 ± 5.000 × 10⁻⁶). No statistically significant difference was observed between PFOS and GenX. However, the P+G group showed significantly lower Complex III activity than PFOS alone (****p* = 0.0008) and GenX alone (*****p* < 0.0001) (Fig. 1E).

At 75 ppm, PFOS (0.6067 ± 0.05196, *****p* = 0.0002), GenX (0.6068 ± 0.02173, ****p* = 0.0004), and P+G (0.3142 ± 0.06653, *****p* < 0.0001) significantly reduced Complex III activity compared with the control (1.000 ± 5.000 × 10⁻⁶). No statistically significant difference was observed between PFOS and GenX. However, Complex III activity in the P+G group was significantly lower than in the PFOS (***p* = 0.0055) and GenX (***p* = 0.0082) groups (Fig. 1F).

The specificity of Complex III activity was assessed using antimycin A, a selective Complex III inhibitor. Antimycin A significantly reduced activity to 0.2370 ± 0.03538 (*****p* < 0.0001) compared with the control (Fig. 1G). Complex-specific inhibitors were used during each assay run to assess the specificity of the measured enzyme activities.

In addition to Complex III, we examined the effects of PFOS and GenX on mitochondrial Complex IV activity in isolated mouse liver mitochondria following a 15-min exposure. PFOS significantly reduced Complex IV activity at all concentrations tested, with normalized activity decreasing from 1.000 ± 7.000 × 10⁻⁶ in the control group to 0.6750 ± 0.04042 at 25 ppm (*****p* < 0.0001), 0.5779 ± 0.04849 at 50 ppm (*****p* < 0.0001), and 0.5687 ± 0.06367 at 75 ppm (*****p* < 0.0001) (Fig. 1H). In contrast, GenX did not significantly alter Complex IV activity at 25 ppm (0.8965 ± 0.03276, *p* > 0.05) or 50 ppm (0.8768 ± 0.02466, *p* > 0.05). However, Complex IV activity was significantly reduced at 75 ppm (0.7316 ± 0.04556, ***p* = 0.0047) compared with the control (Fig. 1I).

Combined exposure to PFOS and GenX (P+G) significantly reduced Complex IV activity at all concentrations tested. Complex IV activity decreased from 1.000 ± 7.000 × 10⁻⁶ in the control group to 0.3262 ± 0.05439 at 25 ppm (*****p* < 0.0001), 0.2604 ± 0.02522 at 50 ppm (*****p* < 0.0001), and 0.2364 ± 0.01948 at 75 ppm (*****p* < 0.0001) (Fig. 1J).

To further compare the effects of individual and combined treatments, Complex IV activity was analyzed at each concentration (Fig. 1K–M). At 25 ppm, PFOS significantly reduced Complex IV activity to 0.6750 ± 0.04042 (*****p* < 0.0001), whereas GenX did not significantly alter activity (0.8965 ± 0.03276, *p* > 0.05), compared with the control (1.000 ± 7.000 × 10⁻⁶). Combined exposure resulted in a greater reduction in Complex IV activity (0.3262 ± 0.05439, *****p* < 0.0001) compared with the control. PFOS also showed significantly lower Complex IV activity than GenX (***p* = 0.0056). The P+G group showed significantly lower activity than both PFOS (*****p* < 0.0001) and GenX (*****p* < 0.0001) alone (Fig. 1K).

At 50 ppm, PFOS significantly reduced Complex IV activity to 0.5779 ± 0.04849 (*****p* < 0.0001), whereas GenX showed a reduction that was not statistically significant (0.8768 ± 0.02466, *p* = 0.0779), compared with the control (1.000 ± 5.000 × 10⁻⁶). Combined PFOS and GenX exposure further reduced Complex IV activity to 0.2604 ± 0.02522 (*****p* < 0.0001). PFOS produced significantly greater inhibition than GenX (*****p* < 0.0001), while the P+G group showed significantly lower Complex IV activity than either PFOS (*****p* < 0.0001) or GenX (*****p* < 0.0001) alone (Fig. 1L).

At 75 ppm, Complex IV activity was significantly reduced by PFOS (0.5687 ± 0.06367, *****p* < 0.0001), GenX (0.7316 ± 0.04556, ***p* = 0.0042), and P+G (0.2364 ± 0.01948, *****p* < 0.0001), compared with the control (1.000 ± 7.000 × 10⁻⁶). No statistically significant difference was observed between PFOS and GenX. However, the P+G group showed significantly lower Complex IV activity than PFOS (****p* = 0.0006) and GenX (*****p* < 0.0001) alone (Fig. 1M). The specificity of Complex IV activity was assessed using potassium cyanide (KCN), a Complex IV inhibitor. KCN significantly reduced Complex IV activity to 0.02536 ± 0.001151 (*****p* < 0.0001) compared with the control (Fig. 1N).

PFOS significantly reduced Complex III activity at all concentrations tested, with activity decreasing from 1.000 ± 3.333 × 10⁻⁷ in the control group to 0.8014 ± 0.05247 at 25 ppm (***p* = 0.0068), 0.7372 ± 0.01068 at 50 ppm (****p* = 0.0006), and 0.6623 ± 0.05018 at 75 ppm (*****p* < 0.0001) (Fig. 2A). GenX did not significantly affect Complex III activity at 25 ppm (1.007 ± 0.02020, p > 0.05), but significantly reduced activity at 50 ppm (0.8923 ± 0.06272, **p* = 0.0163) and 75 ppm (0.8700 ± 0.04848, ***p* = 0.0013) (Fig. 2B). Combined PFOS and GenX exposure also significantly reduced Complex III activity at all concentrations, with activity decreasing from 1.000 ± 3.500 × 10⁻⁶ in the control group to 0.7670 ± 0.06585 at 25 ppm (***p* = 0.0042), 0.6099 ± 0.06701 at 50 ppm (*****p* < 0.0001), and 0.4408 ± 0.06514 at 75 ppm (*****p* < 0.0001) (Fig. 2C).

To compare the effects of individual and combined treatments, Complex III activity was further analyzed at each concentration. At 25 ppm, PFOS significantly reduced Complex III activity compared with GenX (**p* = 0.0162). Similarly, the P+G group showed significantly lower activity than GenX (***p* = 0.0051). However, no statistically significant difference was observed between PFOS and P+G (p > 0.05) (Fig. 2D). Both PFOS and P+G significantly reduced Complex III activity compared with the control, whereas GenX did not. At 50 ppm, PFOS significantly reduced Complex III activity compared with GenX (**p* = 0.0144). The P+G group showed significantly lower activity than both PFOS (**p* = 0.0496) and GenX (*****p* < 0.0001) (Fig. 2E). At 75 ppm, PFOS also showed significantly lower Complex III activity than GenX (***p* = 0.0012). Combined exposure further reduced Complex III activity compared with PFOS (****p* = 0.0006) and GenX (*****p* < 0.0001) alone (Fig. 2F). The specificity of Complex III activity was assessed using antimycin A, a selective Complex III inhibitor. Antimycin A significantly reduced Complex III activity to 0.08632 ± 0.01732 (*****p* < 0.0001) compared with the control (Fig. 2G).

We next examined Complex IV activity in isolated brain mitochondria following acute PFOS, GenX, and P+G exposure. PFOS significantly reduced Complex IV activity at all concentrations tested, with activity decreasing from 1.000 ± 1.667 × 10⁻⁷ in the control group to 0.8884 ± 0.02027 at 25 ppm (***p* = 0.0052), 0.8723 ± 0.02374 at 50 ppm (***p* = 0.0017), and 0.8613 ± 0.001016 at 75 ppm (****p* = 0.0008) (Fig. 2H). Similarly, GenX significantly reduced Complex IV activity at 25 ppm (0.8183 ± 0.02402, ***p* = 0.0040), 50 ppm (0.8156 ± 0.01520, ***p* = 0.0035), and 75 ppm (0.7813 ± 0.02702, ****p* = 0.0008) compared with the control group (Fig. 2I). Combined PFOS and GenX exposure also significantly reduced Complex IV activity at 25 ppm (0.7696 ± 0.04201, *****p* < 0.0001), 50 ppm (0.7437 ± 0.001727, *****p* < 0.0001), and 75 ppm (0.6296 ± 0.02633, *****p* < 0.0001) compared with the control group (Fig. 2J).

At 25 ppm, no statistically significant difference in Complex IV activity was observed between PFOS and GenX (*p* > 0.05) or between GenX and P+G (*p* > 0.05). However, the P+G group showed significantly lower Complex IV activity than PFOS alone (**p* = 0.0261) (Fig. 2K). At 50 ppm, combined exposure also significantly reduced Complex IV activity compared with PFOS alone (***p* = 0.0031). No statistically significant differences were observed between PFOS and GenX (*p* > 0.05) or between GenX and P+G (*p* > 0.05) (Fig. 2L). At 75 ppm, the P+G group showed significantly lower Complex IV activity than PFOS (****p* = 0.0004) and GenX (***p* = 0.00154) alone. However, no statistically significant difference was observed between PFOS and GenX (*p* > 0.05) (Fig. 2M). The specificity of Complex IV activity was assessed using potassium cyanide (KCN), a Complex IV inhibitor. KCN significantly reduced Complex IV activity to 0.2251 ± 0.01780 (*****p* < 0.0001) compared with the control (Fig. 2N).

**S2. Repeated exposure: mitochondrial complex activities**

At 0.2 mg/kg BW for 1 month, Complex I activity was significantly increased in the P+G group (1.36023 ± 0.06405, **p* = 0.0391) compared with the control (0.93347 ± 0.09922). Although Complex I activity was also higher in the PFOS (1.18803 ± 0.13501) and GenX (1.15447 ± 0.15879) groups, these differences were not statistically significant (Fig. 3A). No statistically significant reductions were detected for Complex II, III, or IV. Complex II activities were 0.88664 ± 0.15071, 0.74811 ± 0.03871, 0.97629 ± 0.15465, and 0.92436 ± 0.10041 in the control, PFOS, GenX, and P+G groups, respectively. Complex III activities were 0.86586 ± 0.16754, 0.57619 ± 0.06542, 0.86055 ± 0.24922, and 0.67291 ± 0.03174, respectively. Complex IV activities were 0.89765 ± 0.11046, 0.84073 ± 0.07031, 1.06572 ± 0.17102, and 1.04581 ± 0.11927, respectively (Fig. 3A).

At 2 mg/kg BW for 1 month, mitochondrial enzyme activities were significantly reduced in several treatment groups (Fig. 3B). Complex I activity was significantly lower in the GenX (0.56290 ± 0.03215, ***p* = 0.0017) and P+G (0.53221 ± 0.05354, ****p* = 0.0007) groups compared with the control (1.00000 ± 0.07834), whereas PFOS did not significantly alter activity (1.08138 ± 0.15123). Complex II activity was also significantly reduced in the GenX (0.45851 ± 0.04739, *****p* < 0.0001) and P+G (0.46395 ± 0.09477, ****p* = 0.0001) groups compared with the control (1.00000 ± 0.14639). PFOS did not significantly affect Complex II activity (0.87565 ± 0.07969). Complex III activity was significantly reduced in all treatment groups, with activities of 0.66203 ± 0.11197 in PFOS (**p* = 0.0189), 0.47134 ± 0.04276 in GenX (****p* = 0.0001), and 0.35428 ± 0.06809 in P+G (*****p* < 0.0001), compared with the control (1.00000 ± 0.11626). Similarly, Complex IV activity was significantly lower in the GenX (0.46820 ± 0.02932, ****p* = 0.0001) and P+G (0.44918 ± 0.07642, *****p* < 0.0001) groups compared with the control (1.00000 ± 0.03430). No statistically significant difference was observed in the PFOS group (0.90263 ± 0.06857).

At 10 mg/kg BW for 1 month, all three treatments significantly reduced Complex I–IV activities compared with the control (Fig. 3C). Complex I activities were 0.97883 ± 0.04980 in the control, 0.69712 ± 0.05005 in PFOS (****p* = 0.0007), 0.43731 ± 0.07944 in GenX (*****p* < 0.0001), and 0.51332 ± 0.04873 in P+G (*****p* < 0.0001). Complex II activity decreased from 0.99430 ± 0.02158 in the control group to 0.48259 ± 0.06783 in PFOS, 0.18367 ± 0.03194 in GenX, and 0.32167 ± 0.01963 in P+G (all groups, *****p* < 0.0001). Complex III activity was also reduced from 1.00000 ± 0.09608 in the control group to 0.63758 ± 0.04217 in PFOS, 0.34267 ± 0.03524 in GenX, and 0.42295 ± 0.01415 in P+G (all groups, *****p* < 0.0001). Complex IV activity decreased from 1.00926 ± 0.03318 in the control group to 0.54591 ± 0.02833 in PFOS, 0.34383 ± 0.09562 in GenX, and 0.34489 ± 0.04371 in P+G (all groups, *****p* < 0.0001).

At 0.2 mg/kg BW for 5 months, reductions in liver mitochondrial enzyme activities were observed across several complexes (Fig. 3D). Complex I activity was significantly reduced in the GenX (0.42707 ± 0.05547, **p* = 0.0162) and P+G (0.48607 ± 0.02866, **p* = 0.0242) groups compared with the control (1.00000 ± 0.28624), whereas the reduction in the PFOS group (0.56931 ± 0.02733) was not statistically significant. Complex II activity was significantly reduced in all treatment groups, with activities of 0.43420 ± 0.03228 in PFOS (**p* = 0.0178), 0.35812 ± 0.06186 in GenX (***p* = 0.0063), and 0.45083 ± 0.01484 in P+G (**p* = 0.0149), compared with the control (1.00000 ± 0.35179). Complex III activity was significantly reduced only in the GenX group (0.42971 ± 0.05106, **p* = 0.0167) compared with the control (1.00000 ± 0.23959). PFOS (0.57016 ± 0.04731) and P+G (0.63488 ± 0.02884) showed lower mean activities, but these differences were not statistically significant. Complex IV activity was significantly reduced in the GenX (0.45726 ± 0.06748, **p* = 0.0239) and P+G (0.53608 ± 0.01643, **p* = 0.0464) groups compared with the control (1.00000 ± 0.24417). The reduction in the PFOS group (0.60561 ± 0.03877) was not statistically significant.

**Table S1 Mitochondrial respiratory Complex I–IV enzyme activities in isolated mouse liver mitochondria following sub-chronic and chronic PFOS and GenX exposure.**

| Liver | **Control** | | **PFOS 0.2 mg/kg 1-month** | | **GenX 0.2 mg/kg 1-month** | | **P+G 0.2 mg/kg 1-month** | |
| --- | --- | --- | --- | --- | --- | --- | --- | --- |
|  | Mean | SEM | Mean | SEM | Mean | SEM | Mean | SEM |
| C1/CS | 0.9335 | 0.0992 | 1.1880 | 0.1350 | 1.1545 | 0.1588 | 1.3602 | 0.0641 |
| C2/CS | 0.8866 | 0.1507 | 0.7481 | 0.0387 | 0.9763 | 0.1547 | 0.9244 | 0.1004 |
| C3/CS | 0.8659 | 0.1675 | 0.5762 | 0.0654 | 0.8606 | 0.2492 | 0.6729 | 0.0317 |
| C4/CS | 0.8977 | 0.1105 | 0.8407 | 0.0703 | 1.0657 | 0.1710 | 1.0458 | 0.1193 |
| Liver | **Control** | | **PFOS 2 mg/kg 1-month** | | **GenX 2 mg/kg 1-month** | | **P+G 2 mg/kg 1-month** | |
|  | Mean | SEM | Mean | SEM | Mean | SEM | Mean | SEM |
| C1/CS | 1.0000 | 0.0783 | 1.0814 | 0.1512 | 0.5629 | 0.0322 | 0.5322 | 0.0535 |
| C2/CS | 1.0000 | 0.1464 | 0.8757 | 0.0797 | 0.4585 | 0.0474 | 0.4640 | 0.0948 |
| C3/CS | 1.0000 | 0.1163 | 0.6620 | 0.1120 | 0.4713 | 0.0428 | 0.3543 | 0.0681 |
| C4/CS | 1.0000 | 0.0343 | 0.9026 | 0.0686 | 0.4682 | 0.0293 | 0.4492 | 0.0764 |
| Liver | **Control** | | **PFOS 10 mg/kg 1-month** | | **GenX 10 mg/kg 1-month** | | **P+G 10 mg/kg 1-month** | |
|  | Mean | SEM | Mean | SEM | Mean | SEM | Mean | SEM |
| C1/CS | 0.9788 | 0.0498 | 0.6971 | 0.0501 | 0.4373 | 0.0794 | 0.5133 | 0.0487 |
| C2/CS | 0.9943 | 0.0216 | 0.4826 | 0.0678 | 0.1837 | 0.0319 | 0.3217 | 0.0196 |
| C3/CS | 1.0000 | 0.0961 | 0.6376 | 0.0422 | 0.3427 | 0.0352 | 0.4230 | 0.0142 |
| C4/CS | 1.0093 | 0.0332 | 0.5459 | 0.0283 | 0.3438 | 0.0956 | 0.3449 | 0.0437 |
| Liver | **Control** | | **PFOS 0.2 mg/kg 5-month** | | **GenX 0.2 mg/kg 5-month** | | **P+G 0.2 mg/kg 5-month** | |
|  | Mean | SEM | Mean | SEM | Mean | SEM | Mean | SEM |
| C1/CS | 1.0000 | 0.2862 | 0.5693 | 0.0273 | 0.4271 | 0.0555 | 0.4861 | 0.0287 |
| C2/CS | 1.0000 | 0.3518 | 0.4342 | 0.0323 | 0.3581 | 0.0619 | 0.4508 | 0.0148 |
| C3/CS | 1.0000 | 0.2396 | 0.5702 | 0.0473 | 0.4297 | 0.0511 | 0.6349 | 0.0288 |
| C4/CS | 1.0000 | 0.2442 | 0.6056 | 0.0388 | 0.4573 | 0.0675 | 0.5361 | 0.0164 |

After 1 month of exposure at 0.2 mg/kg BW, no statistically significant differences were observed in Complex I–IV activities among the treatment groups (Fig. 4A). Complex I activities were 1.00000 ± 0.05370, 0.78503 ± 0.09472, 0.78732 ± 0.02119, and 0.86086 ± 0.03898 in the control, PFOS, GenX, and P+G groups, respectively. Complex II activities were 1.00000 ± 0.00465, 0.97891 ± 0.10394, 0.77628 ± 0.09481, and 0.95490 ± 0.09108, respectively. Complex III activities were 1.00000 ± 0.04658, 1.00303 ± 0.05535, 0.96426 ± 0.04498, and 1.06566 ± 0.05259, respectively. Complex IV activities were 1.00000 ± 0.04010, 0.95569 ± 0.04860, 0.95523 ± 0.08918, and 1.02925 ± 0.07876, respectively.

Similarly, no statistically significant differences in Complex I–IV activities were detected following 1 month of exposure at 2 mg/kg BW (Fig. 4B). Complex I activities were 1.00000 ± 0.09400, 0.95659 ± 0.10082, 0.78032 ± 0.06441, and 0.79910 ± 0.06446 in the control, PFOS, GenX, and P+G groups, respectively. Complex II activities were 1.00000 ± 0.04956, 1.01784 ± 0.05640, 0.97511 ± 0.06597, and 1.03777 ± 0.08999, respectively. Complex III activities were 1.00000 ± 0.04227, 1.13740 ± 0.14176, 0.98195 ± 0.08458, and 0.95745 ± 0.07495, respectively. Complex IV activities were 1.00000 ± 0.06116, 1.04719 ± 0.08735, 1.02926 ± 0.04440, and 1.10642 ± 0.08690, respectively.

At 10 mg/kg BW for 1 month, Complex I–IV activities also remained statistically unchanged across the treatment groups (Fig. 4C). Complex I activities were 1.00000 ± 0.12434, 1.00546 ± 0.15706, 0.94445 ± 0.02948, and 0.90317 ± 0.05049 in the control, PFOS, GenX, and P+G groups, respectively. Complex II activities were 1.00000 ± 0.18278, 0.92577 ± 0.03480, 0.97474 ± 0.02931, and 1.19731 ± 0.04631, respectively. Complex III activities were 1.00000 ± 0.07635, 0.89516 ± 0.04421, 0.80810 ± 0.09310, and 1.03130 ± 0.11446, respectively. Complex IV activities were 1.00000 ± 0.15772, 0.78790 ± 0.05948, 0.75414 ± 0.09763, and 1.17220 ± 0.09022, respectively.

At 0.2 mg/kg BW for 5 months, several changes in brain mitochondrial enzyme activities were observed (Fig. 4D). Complex I activities were 1.00000 ± 0.03609 in the control group, 1.42052 ± 0.04783 in PFOS (****p* = 0.0003), 1.17356 ± 0.05519 in GenX, and 1.30209 ± 0.15252 in P+G (**p* = 0.0151). Complex I activity was significantly higher in the PFOS and P+G groups compared with the control, whereas the difference in the GenX group was not statistically significant. No significant differences were reported among the PFOS, GenX, and P+G groups. Complex II activity was significantly reduced only in the GenX group (0.71479 ± 0.04566, **p* = 0.0242) compared with the control (1.00000 ± 0.07088). PFOS (1.09095 ± 0.07194) and P+G (0.89918 ± 0.09334) did not significantly alter Complex II activity compared with the control. However, Complex II activity differed significantly between PFOS and GenX (***p* = 0.0015). Complex III activities were 1.00000 ± 0.05273 in the control, 1.36669 ± 0.06996 in PFOS (***p* = 0.0021), 0.93634 ± 0.04461 in GenX, and 0.88723 ± 0.05047 in P+G. Complex III activity was significantly increased in the PFOS group compared with the control, whereas GenX and P+G did not significantly alter activity. Complex III activity was also significantly different between PFOS and GenX (****p* = 0.0002) and between PFOS and P+G (*****p* < 0.0001). No statistically significant differences in Complex IV activity were observed between the control and treatment groups or among the individual and combined treatments. Complex IV activities were 1.00000 ± 0.02206 in the control, 1.05277 ± 0.06266 in PFOS, 0.94157 ± 0.04286 in GenX, and 0.95277 ± 0.08273 in P+G.

**Table S2 Mitochondrial respiratory Complex I–IV enzyme activities in isolated mouse brain mitochondria following sub-chronic and chronic PFOS and GenX exposure.**

| Brain | **Control** | | **PFOS 0.2 mg/kg 1-month** | | **GenX 0.2 mg/kg 1-month** | | **P+G 0.2 mg/kg 1-month** | |
| --- | --- | --- | --- | --- | --- | --- | --- | --- |
|  | Mean | SEM | Mean | SEM | Mean | SEM | Mean | SEM |
| C1/CS | 1.0000 | 0.0537 | 0.7850 | 0.0947 | 0.7873 | 0.0212 | 0.8609 | 0.0390 |
| C2/CS | 1.0000 | 0.0047 | 0.9789 | 0.1039 | 0.7763 | 0.0948 | 0.9549 | 0.0911 |
| C3/CS | 1.0000 | 0.0466 | 1.0030 | 0.0554 | 0.9643 | 0.0450 | 1.0657 | 0.0526 |
| C4/CS | 1.0000 | 0.0401 | 0.9557 | 0.0486 | 0.9552 | 0.0892 | 1.0293 | 0.0788 |
| Brain | **Control** | | **PFOS 2 mg/kg 1-month** | | **GenX 2 mg/kg 1-month** | | **P+G 2 mg/kg 1-month** | |
|  | Mean | SEM | Mean | SEM | Mean | SEM | Mean | SEM |
| C1/CS | 1.0000 | 0.0940 | 0.9566 | 0.1008 | 0.7803 | 0.0644 | 0.7991 | 0.0645 |
| C2/CS | 1.0000 | 0.0496 | 1.0178 | 0.0564 | 0.9751 | 0.0660 | 1.0378 | 0.0900 |
| C3/CS | 1.0000 | 0.0423 | 1.1374 | 0.1418 | 0.9820 | 0.0846 | 0.9575 | 0.0750 |
| C4/CS | 1.0000 | 0.0612 | 1.0472 | 0.0874 | 1.0293 | 0.0444 | 1.1064 | 0.0869 |
| Brain | **Control** | | **PFOS 10 mg/kg 1-month** | | **GenX 10 mg/kg 1-month** | | **P+G 10 mg/kg 1-month** | |
|  | Mean | SEM | Mean | SEM | Mean | SEM | Mean | SEM |
| C1/CS | 1.0000 | 0.1243 | 1.0055 | 0.1571 | 0.9445 | 0.0295 | 0.9032 | 0.0505 |
| C2/CS | 1.0000 | 0.1828 | 0.9258 | 0.0348 | 0.9747 | 0.0293 | 1.1973 | 0.0463 |
| C3/CS | 1.0000 | 0.0764 | 0.8952 | 0.0442 | 0.8081 | 0.0931 | 1.0313 | 0.1145 |
| C4/CS | 1.0000 | 0.1577 | 0.7879 | 0.0595 | 0.7541 | 0.0976 | 1.1722 | 0.0902 |
| Brain | **Control** | | **PFOS 0.2 mg/kg 5-month** | | **GenX 0.2 mg/kg 5-month** | | **P+G 0.2 mg/kg 5-month** | |
|  | Mean | SEM | Mean | SEM | Mean | SEM | Mean | SEM |
| C1/CS | 1.0000 | 0.0361 | 1.4205 | 0.0478 | 1.1736 | 0.0552 | 1.3021 | 0.1525 |
| C2/CS | 1.0000 | 0.0709 | 1.0910 | 0.0719 | 0.7148 | 0.0457 | 0.8992 | 0.0933 |
| C3/CS | 1.0000 | 0.0527 | 1.3667 | 0.0700 | 0.9363 | 0.0446 | 0.8872 | 0.0505 |
| C4/CS | 1.0000 | 0.0221 | 1.0528 | 0.0627 | 0.9416 | 0.0429 | 0.9528 | 0.0827 |

**S3. Body weight and water intake**

To examine the effects of repeated PFAS exposure on body weight, mice were treated with PFOS, GenX, or their combination (P+G) at 0.2, 2, and 10 mg/kg BW for 1 month. An additional group received 0.2 mg/kg BW for 5 months to examine the effects of prolonged exposure. Body weight and water intake were monitored throughout the exposure period (Fig. S1).

**Sub-chronic exposure for 1 month**

At 0.2 mg/kg BW for 1 month, no statistically significant differences in body weight change were observed among the treatment groups. Body weight increased by 2.16667 ± 0.54263 g in the control group, 2.83333 ± 0.40139 g in the PFOS group (*p* > 0.05), 2.000 ± 0.36515 g in the GenX group (*p* > 0.05), and 2.83333 ± 0.40139 g in the P+G group (*p* > 0.05), compared with the control (Fig. S1A). The time-course data showed a gradual increase in body weight across all groups, with no consistent differences in weight gain throughout the treatment period (Fig. S1B). Similarly, water intake fluctuated during the 1-month exposure period, with no statistically significant differences observed among the control, PFOS, GenX, and P+G groups (Fig. S1C).

At 2 mg/kg BW, mice exposed to P+G showed a significant reduction in body weight change compared with the control group, whereas PFOS and GenX alone did not significantly affect body weight change. Body weight increased by 2.8333 ± 0.79232 g in the control group, 2.5000 ± 0.56273 g in the PFOS group (*p* > 0.05), and 1.5000 ± 0.42817 g in the GenX group (*p* > 0.05). In contrast, the P+G group showed a reduction in body weight of -3.000 ± 0.89443 g (*****p* < 0.0001) compared with the control. The reduction in body weight change in the P+G group was also significant compared with PFOS alone (*****p* < 0.0001) and GenX alone (*****p* < 0.0001) (Fig. S1D). The time-course data also showed a progressive decline in body weight in the P+G group toward the end of treatment, whereas mice in the control, PFOS, and GenX groups generally maintained or gained weight (Fig. S1E). Water intake also differed among treatment groups at 2 mg/kg BW (Fig. S1F). Both GenX and P+G groups showed significantly higher water intake than the control group (*****p* < 0.0001 for GenX and *****p* < 0.0001 for P+G).

At 10 mg/kg BW, all treatment groups showed significantly lower body weight changes than the control group. The control group gained 2.833 ± 0.833 g, whereas the PFOS, GenX, and P+G groups showed body weight changes of -5.167 ± 0.654 g (*****p* < 0.0001), -2.667 ± 0.667 g (*****p* < 0.0001), and -8.000 ± 1.238 g (*****p* < 0.0001), respectively, compared with the control (Fig. S1G). Among the treatment groups, PFOS caused a significantly greater reduction in body weight than GenX (**p* = 0.0342). The P+G group showed significantly greater body weight loss than both PFOS (**p* = 0.0117) and GenX (*****p* < 0.0001) alone. The time-course data showed a progressive decline in body weight in the P+G group, particularly during the later stages of treatment. Similar declines were observed in the PFOS and GenX groups, although the reduction was greater in the P+G group (Fig. S1H). At 10 mg/kg BW, water intake was significantly lower in the PFOS (**p* = 0.0251) and P+G (*****p* < 0.0001) groups compared with the control (Fig. S1I).

**Chronic exposure at 0.2 mg/kg BW for 5 months**

To examine the effects of prolonged low-dose PFAS exposure, mice were treated with PFOS, GenX, or their combination (P+G) at 0.2 mg/kg BW for 5 months. Body weight increased in all groups throughout the treatment period, although the P+G group showed less weight gain toward the end of exposure.

At the end of the 5-month treatment, body weight increased by 12.20000 ± 1.48922 g in the control group, 13.00000 ± 1.57762 g in the PFOS group, 10.90000 ± 1.32874 g in the GenX group, and 7.50000 ± 1.18556 g in the P+G group. Compared with the control, body weight gain was significantly lower in the P+G group (***p* = 0.0012), whereas no statistically significant differences were observed in the PFOS or GenX groups. The P+G group also showed significantly lower body weight gain than PFOS alone (*****p* < 0.0001) and GenX alone (**p* = 0.0357). No significant difference was observed between PFOS and GenX (Fig. S1J).

Absolute body weight gradually increased in all treatment groups throughout the 5-month exposure period (Fig. S1K). At the beginning of treatment, average body weights were 29.50000 ± 0.56273 g in the control group, 29.60000 ± 0.58119 g in the PFOS group, 29.70000 ± 0.53852 g in the GenX group, and 30.20000 ± 0.35901 g in the P+G group. At the end of treatment, average body weights increased to 41.70000 ± 1.59199 g in the control group, 42.60000 ± 1.37598 g in the PFOS group, 40.60000 ± 1.29271 g in the GenX group, and 37.70000 ± 1.13578 g in the P+G group. Although body weight increased in all groups, the P+G group showed the lowest mean body weight at the end of treatment. This pattern was consistent with the significantly lower body weight gain observed in the P+G group compared with the control and individual treatment groups (Fig. S1J–K). Water intake fluctuated throughout the 5-month exposure period, with no consistent differences observed among the control, PFOS, GenX, and P+G groups (Fig. S1L).

These findings show that chronic combined PFOS and GenX exposure significantly reduced body weight gain at 0.2 mg/kg BW, although mice in all groups continued to gain weight throughout treatment. In contrast, no significant differences in body weight change were detected following 1 month of exposure at the same administered dose. This difference suggests that the effects of combined PFAS exposure on body weight may become more apparent with chronic treatment. However, direct statistical comparisons between the two exposure durations are needed to determine whether body weight responses differ significantly.

**Figure S1**
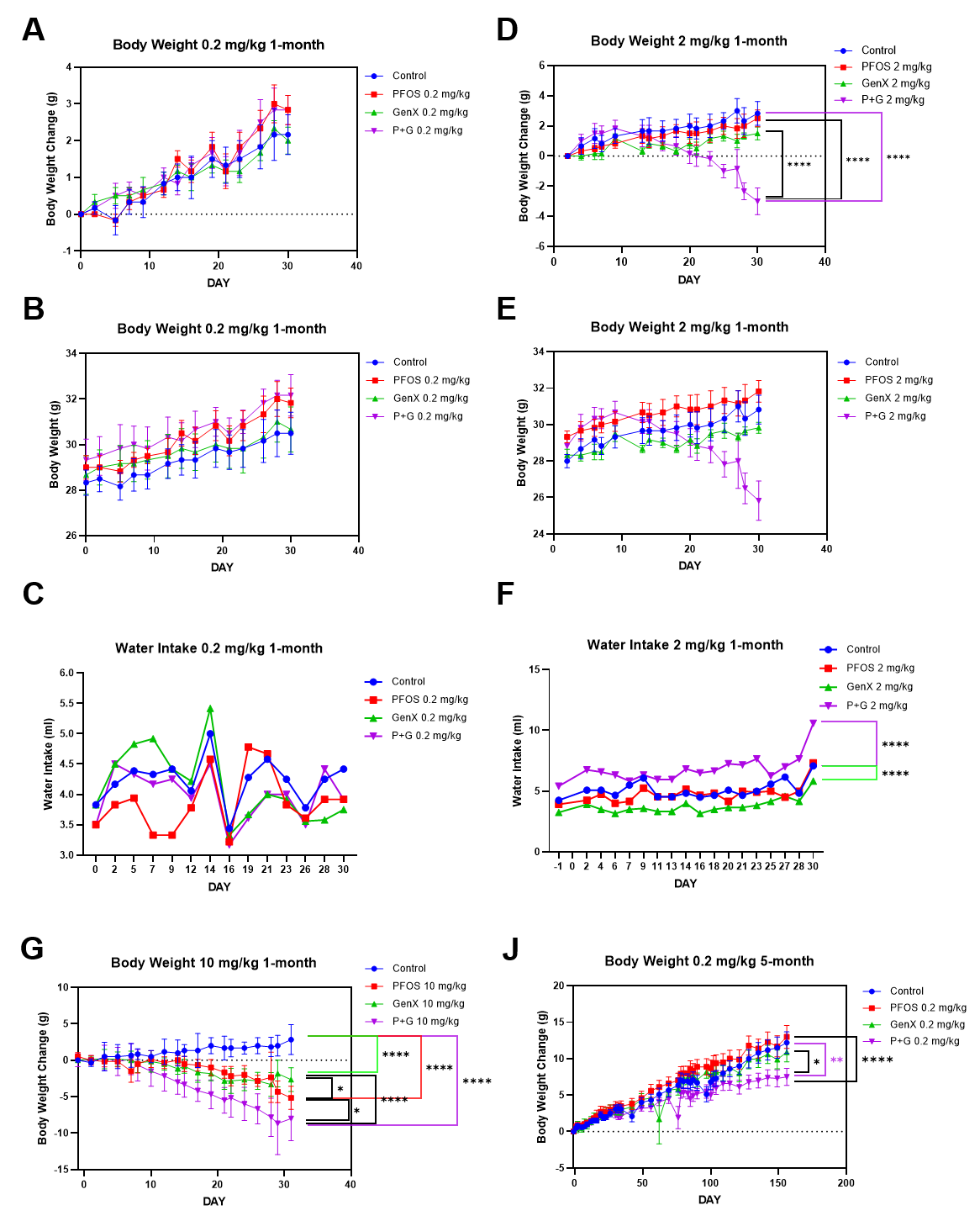

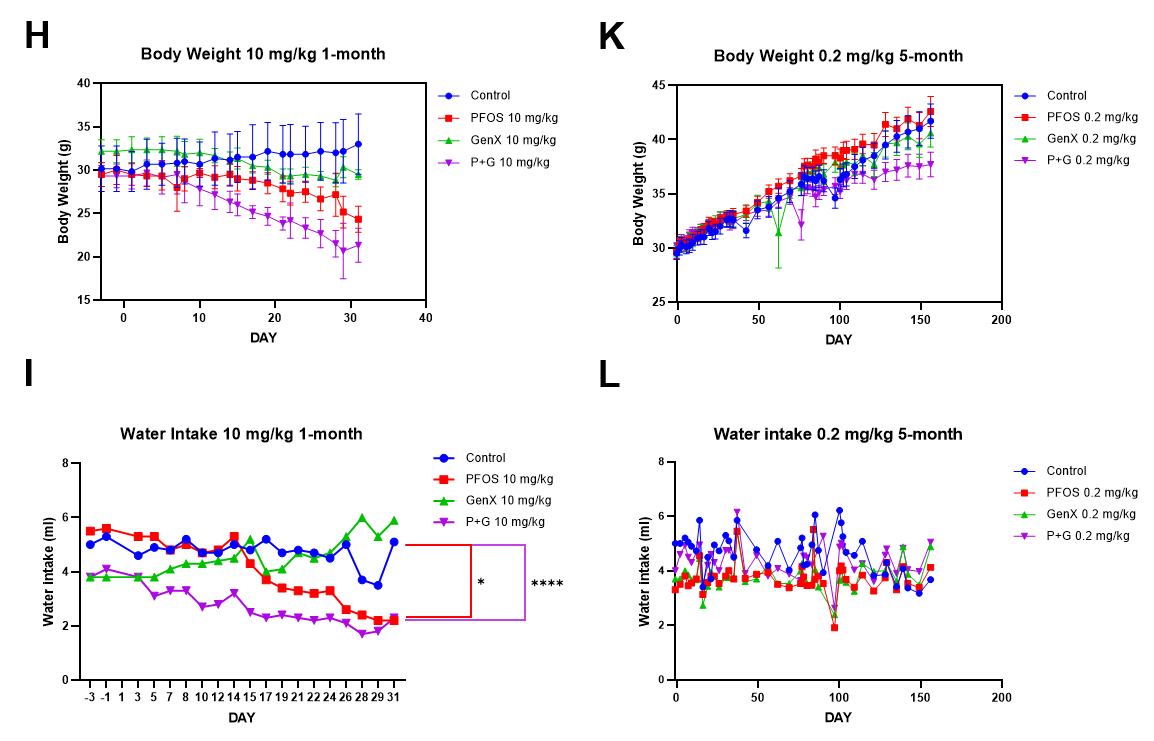
**Figure S1. Effects of sub-chronic and chronic PFOS and GenX exposure on body weight change, absolute body weight, and water intake in mice.**

Mice were exposed to PFOS, GenX, or their combination (P+G) at 0.2, 2, and 10 mg/kg BW for 1 month or at 0.2 mg/kg BW for 5 months. (A–C) (A) Body weight change from baseline, (B) absolute body weight, and (C) water intake following 1 month of exposure at 0.2 mg/kg BW. (D–F) (D) Body weight change from baseline, (E) absolute body weight, and (F) water intake following 1 month of exposure at 2 mg/kg BW. (G–I) (G) Body weight change from baseline, (H) absolute body weight, and (I) water intake following 1 month of exposure at 10 mg/kg BW. (J–L) (J) Body weight change from baseline, (K) absolute body weight, and (L) water intake following 5 months of exposure at 0.2 mg/kg BW. Data are presented as mean ± SEM. Statistical significance was determined using two-way ANOVA followed by Tukey’s multiple-comparisons test. *p < 0.05, **p < 0.01, ***p < 0.001, ****p < 0.0001; ns, not significant.

**S4. PFOS and GenX in mouse plasma and brain regions: detailed Results**

**PFOS and GenX levels in plasma**

At 0.2 mg/kg BW, PFOS was detected in plasma from the PFOS and P+G groups, with concentrations of 4148.33333 ± 482.01395 ng/mL and 8891.66667 ± 671.95345 ng/mL, respectively. PFOS was not detected in the control or GenX-alone groups. Plasma PFOS concentrations were significantly higher in the P+G group than in the PFOS-alone group (**p* = 0.0061). GenX was detected in the GenX (449.33333 ± 67.35561 ng/mL) and P+G (676.33333 ± 43.94062 ng/mL) groups, whereas it was not detected in the control or PFOS-alone groups. Although the mean GenX concentration was higher in the P+G group, the difference between the GenX and P+G groups was not statistically significant (Fig. 5A-B).

At 2 mg/kg BW, plasma PFOS concentrations were 45048 ± 8002.09573 ng/mL in the PFOS group and 102585.33333 ± 27917.52676 ng/mL in the P+G group. Although the mean PFOS concentration was higher following combined exposure, the difference was not statistically significant. GenX concentrations were 4149.386667 ± 393.639 ng/mL in the GenX group and 4142.12533 ± 195.59789 ng/mL in the P+G group, with no significant difference between the two groups (Fig. 5C-D).

At 10 mg/kg BW, plasma PFOS concentrations increased to 258846.33333 ± 43469.97678 ng/mL in the PFOS group and 190499.33333 ± 21198.66509 ng/mL in the P+G group, with no significant difference between treatments. In contrast, GenX concentrations were 23634.66667 ± 666.24328 ng/mL in the GenX group and 3813 ± 1774.34392 ng/mL in the P+G group, with significantly lower GenX concentrations in the P+G group (*****p* < 0.0001) (Fig. 5E-F).

**PFOS and GenX distribution in different brain regions**

At 0.2 mg/kg BW, PFOS was detected in the cerebral cortex at 3.8 ± 0.80208 ng/mg protein in the PFOS group (***p* = 0.0097) and 6.2 ± 0.91652 ng/mg in the P+G group (****p* = 0.0004) compared with the control. No PFOS was detected in the control or GenX-alone groups. Although the mean PFOS concentration was higher in the P+G group than in the PFOS-alone group, the difference was not statistically significant (*p* = 0.0899). GenX was detected at 0.03333 ± 0.03333 ng/mg in the GenX group (*p* = 0.7182) and 0.16333 ± 0.02906 ng/mg in the P+G group (***p* = 0.0035) compared with the control. GenX concentrations were significantly higher in P+G than in GenX alone (**p* = 0.0135) (Fig. 5G-H).

At 2 mg/kg BW, PFOS concentrations in the cerebral cortex were 25.86835 ± 1.69548 in the PFOS group (*p* = 0.0798) and 42.99241 ± 12.62971 ng/mg in the P+G group (***p* = 0.0061) compared with the control. Although the mean PFOS concentration was higher in the P+G group, no significant difference was observed between PFOS and P+G. GenX concentrations were 0.49727 ± 0.15957 ng/mg in the GenX group (**p* = 0.0202) and 0.51820 ± 0.08826 ng/mg in the P+G group (**p* = 0.0162) compared with the control, with no significant difference between the two treatments. (Fig. 5I-J).

In the cerebellum at 2 mg/kg BW, PFOS concentrations were 217.42676 ± 62.71185 ng/mg in the PFOS group (**p* = 0.0103) and 141.03953 ± 32.17613 ng/mg in the P+G group (*p* = 0.0846) compared with the control. No statistically significant difference was observed between PFOS and P+G. GenX concentrations were 0.54910 ± 0.18440 in the GenX group and 1.27769 ± 0.70881 ng/mg in the P+G group, without significant differences between treatment groups. (Fig. 5K-L).

In the brainstem at 2 mg/kg BW, PFOS concentrations were 179.50800 ± 20.53911 ng/mg in the PFOS group (***p* = 0.0053) and 91.21319 ± 47.63149 ng/mg in the P+G group (*p* = 0.1369) compared with the control. The difference between PFOS and P+G was not statistically significant (*p* = 0.1529). GenX concentrations were 0.28234 ± 0.10159 ng/mg in the GenX group and 0.72142 ± 0.50189 ng/mg in the P+G group, without significant differences between treatments (Fig. 5M-N).

At 10 mg/kg BW, PFOS concentrations in the cerebral cortex were 123.06667 ± 44.15938 ng/mg in the PFOS group (**p* = 0.0189), 0.75333 ± 0.15962 ng/mg in the GenX group, and 78.2 ± 5.92497 ng/mg in the P+G group (*p* = 0.1378) compared with the control. No significant difference was observed between PFOS and P+G. GenX concentrations were 2.43 ± 0.61539 ng/mg in the GenX group (***p* = 0.0027) and 0.45 ± 0.13051 ng/mg in the P+G group. GenX concentrations were significantly lower in P+G than in GenX alone (***p* = 0.0092) (Fig. 5O-P).

**S5. PFOS and GenX in rat liver, plasma, and brain regions: detailed Results**

**PFOS and GenX levels in the liver**

PFOS was detected in the liver following PFOS and P+G exposure, with concentrations of 118.933 ± 23.965 ng/mg liver weight in the PFOS group (**p* = 0.0192) and 137.567 ± 35.894 ng/mg liver weight in the P+G group (***p* = 0.0086), compared with the control (0.03667 ± 0.011 ng/mg liver weight). The GenX-alone group showed a low PFOS concentration of 0.141 ± 0.051 ng/mg liver weight (Fig. S2A). Although the mean PFOS concentration was higher in the P+G group than in the PFOS-alone group, the difference was not statistically significant.

Similarly, GenX was detected in the liver of GenX-treated rats (0.423 ± 0.026 ng/mg liver weight, **p* = 0.0136) and P+G-treated rats (0.540 ± 0.142 ng/mg liver weight, ***p* = 0.0032), compared with the control (0.000 ± 0.000 ng/mg liver weight). GenX was not detected in the PFOS-alone group (0.000 ± 0.000 ng/mg liver weight). No statistically significant difference was observed between the GenX and P+G groups (Fig. S2B).

These results confirm the presence of both PFOS and GenX in the liver following repeated exposure. Although the combined treatment showed numerically higher concentrations of both compounds than their respective individual treatments, co-exposure did not significantly alter their measured liver concentrations.

**PFOS and GenX levels in plasma**

In plasma, PFOS concentrations were significantly elevated in both the PFOS and P+G groups compared with the control (0.000 ± 0.000 ng/mg protein). PFOS concentrations were 198231.007 ± 5064.608 ng/mg protein in the PFOS group (*****p* < 0.00001) and 164785.739 ± 13217.041 ng/mg protein in the P+G group (*****p* < 0.00001). PFOS was not detected in the GenX-alone group (0.000 ± 0.000 ng/mg protein). Notably, plasma PFOS concentrations were significantly lower in the P+G group than in the PFOS-alone group (**p* = 0.0411) (Fig. S2C). GenX concentrations were 0.000 ± 0.000 ng/mg protein in the control, 320.623 ± 177.083 ng/mg protein in the PFOS group, 1316.484 ± 821.175 ng/mg protein in the GenX group, and 3313.563 ± 2670.327 ng/mg protein in the P+G group. Although the mean GenX concentration was higher in the P+G group than in the GenX-alone group, no statistically significant differences were detected among groups (Fig. S2D).

In contrast to the liver, where neither compound showed a significant difference between individual and combined exposure, plasma PFOS concentrations were significantly lower following combined treatment. Plasma GenX concentrations showed considerable variability, particularly in the P+G group, with no statistically significant differences among treatment groups.

**PFOS and GenX distribution in different brain regions**

PFOS and GenX were also measured in the cerebral cortex, cerebellum, and brainstem to examine their regional distribution following exposure (Fig. S2E–F).

In the cerebral cortex, PFOS concentrations were significantly higher in the PFOS group (65.062 ± 16.159 ng/mg protein, *****p* < 0.0001) and P+G group (39.205 ± 2.293 ng/mg protein, ***p* = 0.0019) compared with the control (0.000 ± 0.000 ng/mg protein). A low PFOS concentration was also measured in the GenX-alone group (0.197 ± 0.067 ng/mg protein). Although the mean PFOS concentration was lower in the P+G group than in the PFOS-alone group, this difference was not statistically significant. GenX concentrations in the cerebral cortex were 0.000 ± 0.000 ng/mg protein in the control, 0.000 ± 0.000 ng/mg protein in the PFOS group, 0.161 ± 0.089 ng/mg protein in the GenX group, and 0.152 ± 0.035 ng/mg protein in the P+G group. Neither GenX nor P+G exposure significantly altered cortical GenX concentrations compared with the control, and no significant difference was observed between the GenX and P+G groups.

In the cerebellum, PFOS concentrations were significantly elevated in the PFOS group (51.853 ± 3.288 ng/mg protein, *****p* < 0.0001) and P+G group (56.074 ± 10.279 ng/mg protein, *****p* < 0.0001) compared with the control (0.000 ± 0.000 ng/mg protein). The GenX-alone group showed a low PFOS concentration of 0.458 ± 0.209 ng/mg protein. No statistically significant difference was observed between the PFOS and P+G groups.

GenX concentrations in the cerebellum were 0.000 ± 0.000 ng/mg protein in the control, 0.000 ± 0.000 ng/mg protein in the PFOS group, 0.214 ± 0.047 ng/mg protein in the GenX group (**p* = 0.0221), and 0.315 ± 0.093 ng/mg protein in the P+G group (****p* = 0.0006), compared with the control. Although the mean GenX concentration was higher following combined exposure, the difference between GenX and P+G was not statistically significant.

In the brainstem, PFOS concentrations were significantly higher in the PFOS group (67.098 ± 6.276 ng/mg protein, *****p* < 0.0001) and P+G group (62.020 ± 10.630 ng/mg protein, *****p* < 0.0001) compared with the control (0.000 ± 0.000 ng/mg protein). The GenX-alone group showed a PFOS concentration of 0.519 ± 0.215 ng/mg protein. No statistically significant difference was observed between the PFOS and P+G groups.

GenX concentrations in the brainstem were 0.000 ± 0.000 ng/mg protein in the control, 0.000 ± 0.000 ng/mg protein in the PFOS group, 0.189 ± 0.020 ng/mg protein in the GenX group (**p* = 0.0498), and 0.295 ± 0.088 ng/mg protein in the P+G group (***p* = 0.0013), compared with the control. No significant difference was observed between the GenX and P+G groups.

The detection of PFOS and GenX in all three brain regions indicates that both compounds reached brain tissue following repeated exposure. PFOS concentrations were substantially higher than GenX concentrations, under the experimental conditions examined. In the PFOS-alone group, the highest average PFOS concentration was observed in the brainstem, followed by the cerebral cortex and cerebellum. In contrast, the P+G group showed the highest mean PFOS concentration in the brainstem, followed by the cerebellum and cerebral cortex. However, these regional patterns are descriptive, and direct statistical comparisons are required to determine whether PFOS distribution differs significantly among brain regions. GenX was detected at lower concentrations than PFOS in all three brain regions, with significant increases in the cerebellum and brainstem following both GenX and P+G exposure. Although combined exposure resulted in numerically higher GenX concentrations in these two regions, the differences compared with GenX alone were not statistically significant. The presence of both compounds in brain tissue suggests that PFOS and GenX can reach the central nervous system under the experimental conditions used.

**Figure S2**

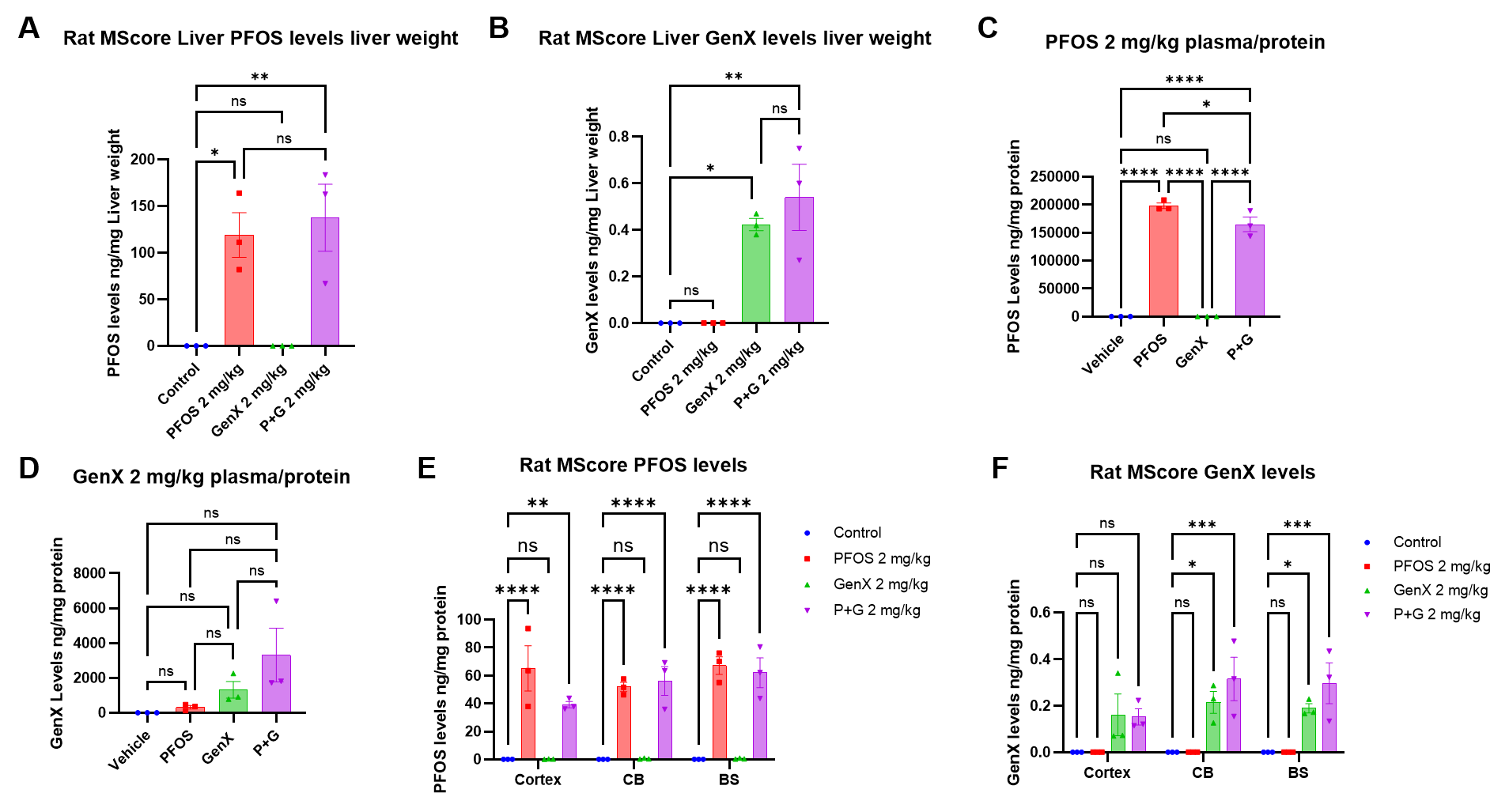
**Figure S2. Distribution of PFOS and GenX in rat liver, plasma, and different brain regions following sub-chronic exposure.**

Rats were exposed to PFOS, GenX, or their combination (P+G) at 2 mg/kg BW. PFOS and GenX concentrations were measured using liquid chromatography–mass spectrometry (LC-MS). (A–B) Concentrations of (A) PFOS and (B) GenX in liver tissues, normalized to liver weight (ng/mg liver weight). (C–D) Plasma concentrations of (C) PFOS and (D) GenX, normalized to protein content (ng/mg protein). (E–F) Concentrations of (E) PFOS and (F) GenX in the cerebral cortex, cerebellum (CB), and brainstem (BS), normalized to protein content (ng/mg protein). Data are presented as mean ± SEM. Individual symbols represent independent biological replicates. Statistical significance was determined using one-way ANOVA followed by Tukey’s multiple-comparisons test for panels A–D and two-way ANOVA followed by Tukey’s multiple-comparisons test for panels E–F. *p < 0.05, **p < 0.01, ***p < 0.001, ****p < 0.0001; ns, not significant.

**S6. OXPHOS protein expression: detailed Results and Western blot methods**

All proteins were quantified using the Pierce™ BCA Protein Assay Kits (23227, Thermo Fisher Scientific Inc., Rockford, IL, USA). Equal amounts of protein (24 µg per samples) were loaded and electrophoresed on 4–20% Mini-PROTEAN TGX Precast Protein Gels (15-well, 8.6 × 6.7 cm; #4561096, Bio-Rad Laboratories, USA) and transferred onto 0.2 µm nitrocellulose membrane (Amersham™ Protran™, 10600001, Cytiva, Germany) using a semi-dry transfer unit.

Membranes were blocked in 5% (w/v) bovine serum albumin (BSA) (BP9706100, Thermo Fisher Scientific, USA) in 0.05% phosphate-buffered saline containing Tween 20 (PBS-T) for 1 hour at room temperature. Membranes were incubated with primary antibodies including total OXPHOS (1:1,000, mouse monoclonal antibody, ab110413, Abcam) and Hsp60 (1:10,000, rabbit monoclonal antibody, ab190828, Abcam) diluted in 0.01% PBS-T for 1 hour at room temperature. Following primary antibody incubation, membranes were incubated with secondary antibodies (1:20,000) including IRDye 680RD goat anti-mouse (#926-68070, LI-COR Biosciences) and IRDye 800CW goat anti-rabbit (#926-32111, LI-COR Biosciences) diluted in 0.01% PBS-T for 2 hours at room temperature. The blots were scanned using a two-channel infrared imaging system (Odyssey Clx, LI-COR Biosciences) and quantified using Image Studio (version 6.1, LI-COR Biosciences).

**Cerebral cortex (2 mg/kg BW for 1 month)**

At 2 mg/kg BW, Complex I protein expression was significantly increased in the PFOS (1.97955 ± 0.263216, *****p* < 0.0001) and GenX (1.714473 ± 0.115262, ****p* = 0.0006) groups compared with the control (1.098789 ± 0.146242), whereas no significant difference was observed in the P+G group (1.010155 ± 0.116443) (Fig. 6A-B). Complex II expression was significantly increased in the GenX group (1.395544 ± 0.094438, **p* = 0.0494) compared with the control (1.0222 ± 0.05514). No significant differences were observed in the PFOS (1.113084 ± 0.123484) or P+G (1.101131 ± 0.053876) groups. In contrast, Complex III expression was significantly reduced in the PFOS group (0.371897 ± 0.100823, ***p* = 0.0039) compared with the control (0.955804 ± 0.044443). No significant differences were observed in the GenX (0.962817 ± 0.035426) or P+G (0.87165 ± 0.051945) groups. Complex IV expression was significantly reduced in the PFOS (0.193305 ± 0.124227, ***p* = 0.0011) and GenX (0.341522 ± 0.102167, **p* = 0.01) groups compared with the control (1 ± 0.118214). Although Complex IV expression was also lower in the P+G group (0.509532 ± 0.172914), this difference was not statistically significant. Complex V expression was significantly increased in the GenX group (1.354712 ± 0.134825, **p* = 0.0467) compared with the control (1 ± 0.053044). No significant differences were observed in the PFOS (1.221296 ± 0.160399) or P+G (0.904048 ± 0.065158) groups. Overall, PFOS and GenX individually altered several OXPHOS proteins in the cerebral cortex, whereas combined exposure did not significantly change any of the five proteins examined.

**Cerebral cortex (10 mg/kg BW for 1 month)**

At 10 mg/kg BW, Complex IV protein expression was significantly increased in all treatment groups compared with the control (1.0000 ± 0.068924). Complex IV expression increased to 1.724346 ± 0.262904 in the PFOS group (***p* = 0.0011), 1.735259 ± 0.152645 in the GenX group (****p* = 0.0009), and 2.086929 ± 0.279973 in the P+G group (*****p* < 0.0001) (Fig. 6C-D). No statistically significant differences were observed in Complex I expression. The levels were 1.120839 ± 0.110994 in the control, 0.946021 ± 0.111897 in PFOS, 0.758369 ± 0.055266 in GenX, and 0.713684 ± 0.034227 in P+G. Similarly, Complex II expression did not differ significantly among treatment groups. The levels were 1.114035 ± 0.074847 in the control, 1.107082 ± 0.128677 in PFOS, 0.713008 ± 0.064438 in GenX, and 0.786988 ± 0.093195 in P+G. No significant differences were observed in Complex III expression, with levels of 1.057949 ± 0.124156 in the control, 0.930592 ± 0.151062 in PFOS, 1.332826 ± 0.147485 in GenX, and 1.146106 ± 0.12605 in P+G. Complex V expression also remained statistically unchanged, with levels of 1 ± 0.075121 in the control, 1.024403 ± 0.131826 in PFOS, 1.021513 ± 0.097086 in GenX, and 0.843326 ± 0.03173 in P+G (Fig. 6C-D). At 10 mg/kg BW, the most prominent change was the significant increase in Complex IV protein expression following all three treatments. Unlike the 2 mg/kg BW groups, where several OXPHOS proteins were altered, significant changes at 10 mg/kg BW were limited to Complex IV.

**Cerebral cortex (0.2 mg/kg BW for 5 months)**

To examine the effects of longer-term PFAS exposure, OXPHOS protein expression was also assessed following treatment at 0.2 mg/kg BW for 5 months.

Complex I expression was significantly reduced in the P+G group (0.562049 ± 0.077869, **p* = 0.0182) compared with the control (1 ± 0.104776). No significant differences were observed in the PFOS (1.029113 ± 0.063899) or GenX (0.819145 ± 0.099275) groups (Fig. 6E-F). Complex II expression was not significantly altered in any treatment group. The levels were 0.974452 ± 0.115009 in the control, 0.926128 ± 0.116124 in PFOS, 1.096781 ± 0.246331 in GenX, and 0.923369 ± 0.114165 in P+G. Similarly, no significant differences were observed in Complex III expression, with levels of 1 ± 0.100094 in the control, 0.981292 ± 0.107338 in PFOS, 0.767496 ± 0.066558 in GenX, and 0.925568 ± 0.060803 in P+G. Complex IV expression was significantly reduced in the GenX (0.587312 ± 0.069724, **p* = 0.0415) and P+G (0.505463 ± 0.132727, **p* = 0.0112) groups compared with the control (1.000 ± 0.122476). Although the mean Complex IV expression was also lower in the PFOS group (0.980083 ± 0.146133), this difference was not statistically significant. No significant differences were observed in Complex V expression. The levels were 1.000 ± 0.09607 in the control, 0.84011 ± 0.064999 in PFOS, 0.81897 ± 0.156647 in GenX, and 0.743375 ± 0.073658 in P+G (Fig. 6E-F).

Following 5 months of exposure at 0.2 mg/kg BW, combined PFOS and GenX exposure significantly reduced Complex I and IV protein expression, whereas GenX alone significantly reduced Complex IV expression. In contrast, PFOS alone did not significantly alter any of the five OXPHOS proteins examined. These results show that chronic exposure at a lower dose was associated with selective changes in mitochondrial OXPHOS protein abundance, particularly following GenX and combined exposure. Across the three exposure conditions, the cerebral cortex showed distinct patterns of OXPHOS protein expression. At 2 mg/kg BW for 1 month, several complexes were affected by PFOS and GenX individually, whereas at 10 mg/kg BW, the significant changes were restricted to increased Complex IV expression. After 5 months at 0.2 mg/kg BW, reduced Complex I and IV expression were observed following combined exposure. These findings suggest that mitochondrial OXPHOS protein expression varies with the compound, exposure level, and duration.

**Cerebellum (2 mg/kg BW for 1 month)**

At 2 mg/kg BW, Complex I protein expression was significantly increased in the GenX (1.295995 ± 0.11631, **p* = 0.0269) and P+G (1.57947 ± 0.130554, *****p* < 0.0001) groups compared with the control (0.971821 ± 0.071667), whereas PFOS did not significantly alter Complex I expression (1.040953 ± 0.066536, p = 0.9022) (Fig. 6G-H). Complex II expression was also significantly increased in the P+G group (1.277808 ± 0.107455, **p* = 0.0127) compared with the control (0.9657 ± 0.027484). No significant differences were observed in the PFOS (0.819684 ± 0.072245) or GenX (0.744177 ± 0.130293) groups. Complex III expression did not differ significantly among treatment groups. The levels were 0.896972 ± 0.043517 in the control, 0.73974 ± 0.064061 in PFOS, 0.775513 ± 0.091928 in GenX, and 0.796402 ± 0.064856 in P+G. In contrast, Complex IV protein expression was significantly reduced in all treatment groups compared with the control (1.080576 ± 0.066361). Complex IV expression decreased to 0.436461 ± 0.079551 in PFOS (*****p* < 0.0001), 0.67004 ± 0.115312 in GenX (***p* = 0.0024), and 0.627148 ± 0.07506 in P+G (***p* = 0.001). No significant differences were observed in Complex V expression, with levels of 1.000 ± 0.031237 in the control, 0.963827 ± 0.051459 in PFOS (p = 0.9792), 1.008293 ± 0.095106 in GenX (p = 0.9997), and 1.13012 ± 0.05291 in P+G (p = 0.5274) (Fig. 6G–H).

**Cerebellum (10 mg/kg BW for 1 month)**

At 10 mg/kg BW, Complex I protein expression was significantly reduced in the PFOS (0.687709 ± 0.127439, ***p* = 0.0064) and P+G (0.581348 ± 0.112948, ****p* = 0.0001) groups compared with the control (1.056596 ± 0.099896). No significant difference was observed in the GenX group (1.022065 ± 0.080464, p = 0.9835) (Fig. 6I-J). Complex II expression was also significantly reduced in the PFOS (0.739798 ± 0.05641, **p* = 0.031) and GenX (0.758587 ± 0.110544, **p* = 0.0226) groups compared with the control (1.056596 ± 0.074524). Although Complex II expression was numerically lower in the P+G group (0.844911 ± 0.113082), the difference was not statistically significant. Complex III protein expression was significantly reduced in the PFOS group (0.728964 ± 0.065484, **p* = 0.043) compared with the control (1 ± 0.050684). Lower mean expression levels were also observed in the GenX (0.747806 ± 0.069487, p = 0.0552) and P+G (0.833535 ± 0.060141, p = 0.2897) groups, but neither difference was statistically significant. Similar to the 2 mg/kg BW exposure, Complex IV protein expression was significantly reduced in all treatment groups. Compared with the control (1.083185 ± 0.058505), Complex IV expression decreased to 0.515285 ± 0.062818 in PFOS (*****p* < 0.0001), 0.594924 ± 0.066969 in GenX (****p* = 0.0002), and 0.509 ± 0.071045 in P+G (*****p* < 0.0001). In contrast, Complex V expression was significantly increased only in the PFOS group (1.324666 ± 0.093439, **p* = 0.0497) compared with the control (1.053012 ± 0.042258). No significant differences were observed in the GenX (1.064188 ± 0.038984) or P+G (1.11895 ± 0.056537) groups.

At 10 mg/kg BW, PFOS exposure significantly reduced Complex I–IV protein expression while increasing Complex V expression. GenX significantly reduced Complex II and IV expression, whereas combined exposure significantly reduced Complex I and IV expression. Compared with the 2 mg/kg BW groups, where Complex I expression increased following GenX and P+G exposure, significant reductions in Complex I expression were observed in the PFOS and P+G groups at 10 mg/kg BW. Although the responses differed among treatments, Complex IV expression was significantly reduced in all treatment groups at both exposure levels.

**Cerebellum (0.2 mg/kg BW for 5 months)**

To examine the effects of prolonged exposure at a lower dose, OXPHOS protein expression was assessed in the cerebellum following 5 months of treatment at 0.2 mg/kg BW. Complex I expression was not significantly altered in any treatment group. The levels were 1 ± 0.090774 in the control, 0.674444 ± 0.056241 in PFOS, 0.89106 ± 0.005127 in GenX, and 1.066114 ± 0.126013 in P+G. Complex II protein expression was significantly increased in the P+G group (1.793704 ± 0.30736, **p* = 0.0106) compared with the control (1.26831 ± 0.034427). No significant differences were observed in the PFOS (1.32777 ± 0.095066) or GenX (1.278067 ± 0.057558) groups. Complex III expression was not significantly altered by any treatment. The levels were 1 ± 0.040711 in the control, 0.933906 ± 0.036173 in PFOS, 1.079173 ± 0.068118 in GenX, and 0.865638 ± 0.09621 in P+G. Similarly, no significant differences were observed in Complex IV expression, with levels of 1 ± 0.094381 in the control, 1.370918 ± 0.198127 in PFOS, 1.218656 ± 0.167845 in GenX, and 0.733683 ± 0.080969 in P+G. Complex V expression also remained statistically unchanged, with levels of 1 ± 0.109009 in the control, 0.923223 ± 0.068427 in PFOS, 1.064402 ± 0.113365 in GenX, and 1.357878 ± 0.169594 in P+G (Fig. 6K-L).

Following 5 months of exposure at 0.2 mg/kg BW, combined PFOS and GenX exposure significantly increased Complex II protein expression, whereas neither compound alone produced significant changes in any of the five OXPHOS proteins examined. Although Complex IV expression was numerically higher in the PFOS and GenX groups and lower in the P+G group, these differences were not statistically significant.

Compared with the 1-month exposure groups, where Complex IV expression was significantly reduced at both 2 and 10 mg/kg BW, no significant change in Complex IV expression was observed after 5 months at 0.2 mg/kg BW. Because both dose and exposure duration differed, these observations describe differences across experimental conditions rather than an established duration-dependent response.

**Brainstem (2 mg/kg BW for 1 month)**

At 2 mg/kg BW, Complex I protein expression was not significantly altered following PFOS, GenX, or P+G treatment compared with the control. In contrast, Complex II protein expression was significantly reduced in the PFOS group (****p* = 0.0004), whereas no significant differences were observed in the GenX or P+G groups. Complex III expression also remained statistically unchanged following PFOS, GenX, and P+G exposure. Complex IV expression was significantly reduced in all treatment groups compared with the control, including PFOS (***p* = 0.0041), GenX (**p* = 0.0445), and P+G (*****p* < 0.0001). No significant differences were observed in Complex V expression following PFOS, GenX, or P+G exposure (Fig. 6M-N).

**Brainstem (10 mg/kg BW for 1 month)**

At 10 mg/kg BW, no statistically significant differences were observed in Complex I–V protein expression following PFOS, GenX, or P+G exposure (Fig. 6O-P).

Complex I expression was 1 ± 0.043147 in the control, 1.001443 ± 0.121205 in PFOS, 1.217146 ± 0.198475 in GenX, and 1.165103 ± 0.174677 in P+G, with no significant differences compared with the control. Similarly, Complex II expression was not significantly altered, with levels of 1 ± 0.138516 in the control, 1.070314 ± 0.133955 in PFOS, 1.031589 ± 0.13859 in GenX, and 1.339998 ± 0.211352 in P+G. Complex III expression remained statistically unchanged, with levels of 1 ± 0.025322 in the control, 0.91609 ± 0.05822 in PFOS, 0.984884 ± 0.132657 in GenX, and 0.930541 ± 0.102051 in P+G. No statistically significant differences were observed in Complex IV expression. The levels were 1 ± 0.189326 in the control, 1.034356 ± 0.114822 in PFOS, 0.982291 ± 0.17252 in GenX, and 1.028235 ± 0.220403 in P+G. Complex V expression also remained statistically unchanged, with levels of 1 ± 0.026467 in the control, 1.094217 ± 0.078329 in PFOS, 1.079136 ± 0.150594 in GenX, and 1.178553 ± 0.170148 in P+G.

**Brainstem (0.2 mg/kg BW for 5 months)**

To examine the effects of longer-term exposure at a lower dose, OXPHOS protein expression was assessed in the brainstem following 5 months of treatment at 0.2 mg/kg BW. No significant differences were observed in Complex I expression. The levels were 1 ± 0.178979 in the control, 0.806828 ± 0.046536 in PFOS, 0.806465 ± 0.084369 in GenX, and 0.817448 ± 0.10718 in P+G. Similarly, Complex II expression was not significantly altered, with levels of 1 ± 0.220378 in the control, 0.732057 ± 0.089287 in PFOS, 0.829075 ± 0.119036 in GenX, and 0.846133 ± 0.087727 in P+G. Complex III expression also remained statistically unchanged, with levels of 1 ± 0.225861 in the control, 1.291596 ± 0.05884 in PFOS, 1.042579 ± 0.089823 in GenX, and 1.080907 ± 0.023612 in P+G.

In contrast, Complex IV protein expression was significantly increased in the P+G group (1.697062 ± 0.256672, ***p* = 0.0022) compared with the control (1.000 ± 0.219861). Although Complex IV expression was also numerically higher in the PFOS (1.383496 ± 0.047677, p = 0.128) and GenX (1.27947 ± 0.101717, p = 0.3388) groups, these differences were not statistically significant. No significant differences were observed in Complex V expression, with levels of 1.0000 ± 0.188865 in the control, 0.882876 ± 0.038843 in PFOS (p = 0.8734), 0.871109 ± 0.077689 in GenX (p = 0.8402), and 0.933256 ± 0.040801 in P+G (p = 0.9718) (Fig. 6Q-R).

Following 5 months of exposure at 0.2 mg/kg BW, combined PFOS and GenX exposure significantly increased Complex IV protein expression, whereas neither compound alone produced significant changes in any of the five OXPHOS subunits examined. The brainstem showed selective alterations in OXPHOS protein expression following PFOS and GenX exposure. At 2 mg/kg BW for 1 month, Complex IV expression was reduced following individual and combined treatments, while Complex II expression was reduced only following PFOS exposure. At 10 mg/kg BW, no significant changes were detected in any of the five OXPHOS proteins. In contrast, chronic exposure at 0.2 mg/kg BW was associated with increased Complex IV expression following combined treatment.

**S7. Molecular docking: methods detail and Table S3**

For Complex III, the Qo and Qi binding sites were identified using experimentally resolved ubiquinone molecules in the human structure. Equivalent mouse sites were identified through structural alignment, which produced a Cα RMSD of 0.676 Å across 378 aligned atoms. For Complex IV, the heme a₃–CuB catalytic region was selected as an initial docking region. The distance between heme a₃ iron and CuB was 4.49 Å in human CIV and 5.16 Å in mouse CIV, and the midpoint between these atoms was used to center the docking box. All calculations were performed using AutoDock Vina v1.2.7, with a rigid receptor, a 24 × 24 × 24 Å search box, 10 requested poses, and random seed 20261027. Exhaustiveness was 32 for CIV, mouse CIII, and human CIII Qi experiments, and for the human CIII Qo mixture; the initial human CIII Qo single-ligand calculations used exhaustiveness 16. PFOS and GenX were docked individually and simultaneously, with the mixture represented by one molecule of each compound.

Table S3. **Predicted docking scores of PFOS, GenX, and their combined configuration with human and mouse mitochondrial Complexes III and IV.**

| **Complex / species** | **Binding region** | **PFOS** | **GenX** | **P+G** |
| --- | --- | --- | --- | --- |
| III / Human | Qo | −8.299 | −6.877 | −11.730 |
| III / Human | Qi | −8.711 | −7.407 | −12.070 |
| III / Mouse | Qo | −7.634 | −6.402 | −10.580 |
| III / Mouse | Qi | −6.882 | −7.084 | −11.440 |
| IV / Human | Heme a₃–CuB | 5.146 | −1.914 | 9.454 |
| IV / Mouse | Heme a₃–CuB | −0.1504 | −5.160 | −0.7048 |
